# Dynamical management of potential threats regulated by dopamine and direct- and indirect-pathway neurons in the tail of the striatum

**DOI:** 10.1101/2022.02.05.479267

**Authors:** Iku Tsutsui-Kimura, Naoshige Uchida, Mitsuko Watabe-Uchida

## Abstract

Avoiding potential threats before experiencing an actual outcome is critical to prevent a disaster. Here we examined roles of the tail of the striatum (TS) and its dopamine input in threat management. Mice were presented with a potential threat (a moving object) while pursuing rewards. Mice initially failed to obtain rewards, but gradually successfully obtained rewards in later trials. We show that the initial failures depended on dopamine and direct-pathway neurons in TS, and variability in failure rate across trials and individuals was positively correlated with the activity of these neurons. In contrast, indirect-pathway neurons in TS were critical for eventual improvement in reward acquisition, and their activity was positively correlated with successful reward acquisition. These results demonstrate that direct- and indirect-pathway TS neurons promote and suppress threat avoidance, respectively, at different stages, providing a mechanism for overcoming a potential threat while maintaining the threat estimates.

## INTRODUCTION

In natural environments, animals consider both rewarding and threatening future outcomes to select appropriate actions for survival. Although rewards and threats can be regarded as opposing factors, critical asymmetry exists (Boureau and Dayan, 2011). While animals often experience actual rewarding outcomes and can learn from such experiences in daily life, a single instance of actual, ultimate outcomes of threats such as pain, injury and death can be catastrophic. It is therefore critical for animals to consider threat potential for action selection to prevent ultimate disastrous outcomes. On the other hand, being too cautious to potential threats would also harmful since animals would not be able to meet their needs. Because too much or too little caution against potential threats would be costly (Blanchard, 1991; Dunsmoor et al., 2015), it is critical to properly assess potential threats to maintain normal life. How the brain estimates potential threats in uncertain environments, however, remains largely unknown.

Threat management requires balancing threat avoidance against other factors (such as reward opportunity) for behavioral choice. When animals overcome a potential threat under threat-reward conflict, it is not practical to erase knowledge of a potential threat too quickly because the threat estimates would be useful in the future. Instead, animals need to temporarily suppress effects of a threat on actions. Thus, overcoming threat can be fundamentally different from extinction, though many fear extinction experiments may have both behavioral components (Dunsmoor et al., 2015). Overcoming a potential threat is critical in many situations in human life as well, and maladaptation to potential threats can cause risky behaviors in adolescence (Steinberg, 2007) and withdrawal in anxiety and post-traumatic stress disorder (Blanchard, 1991; Dunsmoor et al., 2015). However, most previous studies have focused on extinction, rather than specifically targeting overcoming threats. Further, painful stimuli are commonly used in extinction studies (Dunsmoor et al., 2015; Herry et al., 2010; Pare and Duvarci, 2012; Parsons and Ressler, 2013; Sangha et al., 2020; Sotres-Bayon et al., 2006), rather than potential threats, despite the fact that natural threats do not necessarily involve physical pain. It thus remains unclear how the brain can overcome potential threats while preserving threat knowledge.

Dopamine has long been studied in the context of reward-related behaviors (Schultz, 1998). Based on observed dopamine activity patterns and effects of their manipulation, it is well accepted that dopamine broadcasts reward prediction error, the discrepancy between actual and expected reward value, as evaluation signals for animals to update reward prediction and to learn to perform actions that lead to reward (Bayer and Glimcher, 2005; Cohen et al., 2012; Hart et al., 2014; Schultz et al., 1997). In addition, dopamine release can enhance signaling of downstream brain areas, promoting approach to and learning of rewarding stimuli (Kakade and Dayan, 2002; Lahiri and Bevan, 2020; Saunders and Robinson, 2012). In contrast, recent studies found that there is a unique subpopulation of dopamine neurons. Dopamine neurons that project to the tail of the striatum (TS) receive distinct sets of presynaptic inputs (Menegas et al., 2015) and show unique activity patterns; this subpopulation of dopamine neurons signals physical salience of external stimuli in multiple modalities, but does not reliably signal reward values (Kim et al., 2015; Menegas et al., 2018; Menegas et al., 2017). Further distinguishing this population, optogenetic activation of TS-projecting dopamine neurons promotes avoidance, instead of producing the positively reinforcing effect commonly observed in canonical dopamine signaling (Menegas et al., 2018).

Notably, previous studies found that dopamine in TS facilitates avoidance of a novel object (Akiti et al., 2021; Menegas et al., 2018). From behavioral and computational analyses of novelty-induced behaviors, these studies proposed a simple reinforcement learning model where dopamine in TS facilitates avoidance of a potential threat by two mechanisms: First, dopamine in TS estimates potential threats based on the physical salience of a novel stimulus (’shaping bonus’ in reinforcement learning (Kakade and Dayan, 2002)), and second, dopamine in TS is used to develop threat prediction for future behavioral choice (Akiti et al., 2021). As a result, novelty avoidance, facilitated by TS dopamine, may compete with the drive for novelty approach triggered by a separate system, which models action selection under fear-curiosity conflicts. This idea opens up the novel possibility that TS plays a general role in the avoidance of potential threats based on its physical salience under threat-reward conflicts, in a similar manner to novelty exploration.

In the present study, we examined the neural mechanisms underlying threat management, particularly focusing on the role of TS. We modified a recently established behavioral paradigm using a fictive predator (Amir et al., 2015; Choi and Kim, 2010). In this paradigm, a novel threatening object (’monster’ hereafter) was introduced while mice were navigating in a foraging arena to acquire water rewards. Upon the introduction of a monster, mice immediately avoided a moving monster, and quickly learned to avoid approaching it, but then gradually overcame the threat and started harvesting rewards successfully.

Herein, we demonstrate that threat avoidance depended on physical salience (size and movement) of the object. Furthermore, dopamine in TS was activated in response to the monster, and avoidance critically depended on dopamine in TS. We show that specific aspects of threat management were differentially controlled by different cell types in TS: medium spiny neurons (MSNs) in direct and indirect pathways (dMSN and iMSN, respectively), whose circuit architecture suggests opposing functions (DeLong, 1990; Gerfen et al., 1990; Gerfen and Young, 1988). Our results reveal a neural substrate of threat management: iMSNs promote overcoming threat, while dMSNs maintain an estimate of a potential threat that is facilitated by dopamine in TS.

## RESULTS

### Dynamical management of potential threat

To gain insight into the mechanisms underlying threat management, we developed a “monster task” where mice freely forage in an open arena with occasional existence of an unknown moving ’monster’ object (Figure 1). The monster apparatus consists of a shelter and a foraging arena divided by a door (Figure 1A, left). The shelter is small and dark, while the foraging arena is long with a small hole on the ground for liquid reward. In the monster paradigm, each session starts while a mouse stays in a shelter with the door closed. At trial start, the door opens, allowing the mouse to freely go out to the foraging arena. In each trial, only one drop of water is available. After the mouse acquires a drop of water reward, it must return to the shelter to trigger the next trial. Until then, the mouse can freely explore the arena, but no more reward is available. Once the mouse returns to the shelter, the door closes, and the next trial starts after some delay (inter-trial-interval). In some sessions, a ’monster’ object was set at the far end of the foraging arena. The object was attached to a motorized bar so that it could move back and forth along the longitudinal axis of the foraging arena, but never overlapped with the reward hole. In these sessions (which we will call “monster session” hereafter), there is an invisible line which determines the “monster territory”. Once the mouse crosses this line to enter the monster territory, the monster starts to ’charge’, moving back and forth with a tone until the mouse goes back to the shelter. To minimize human interruption, all the task procedures such as monster movement, door closing, and reward delivery are automatically controlled based on the animal’s position monitored using infrared break-beam sensors (see Methods).

**Figure 1.**
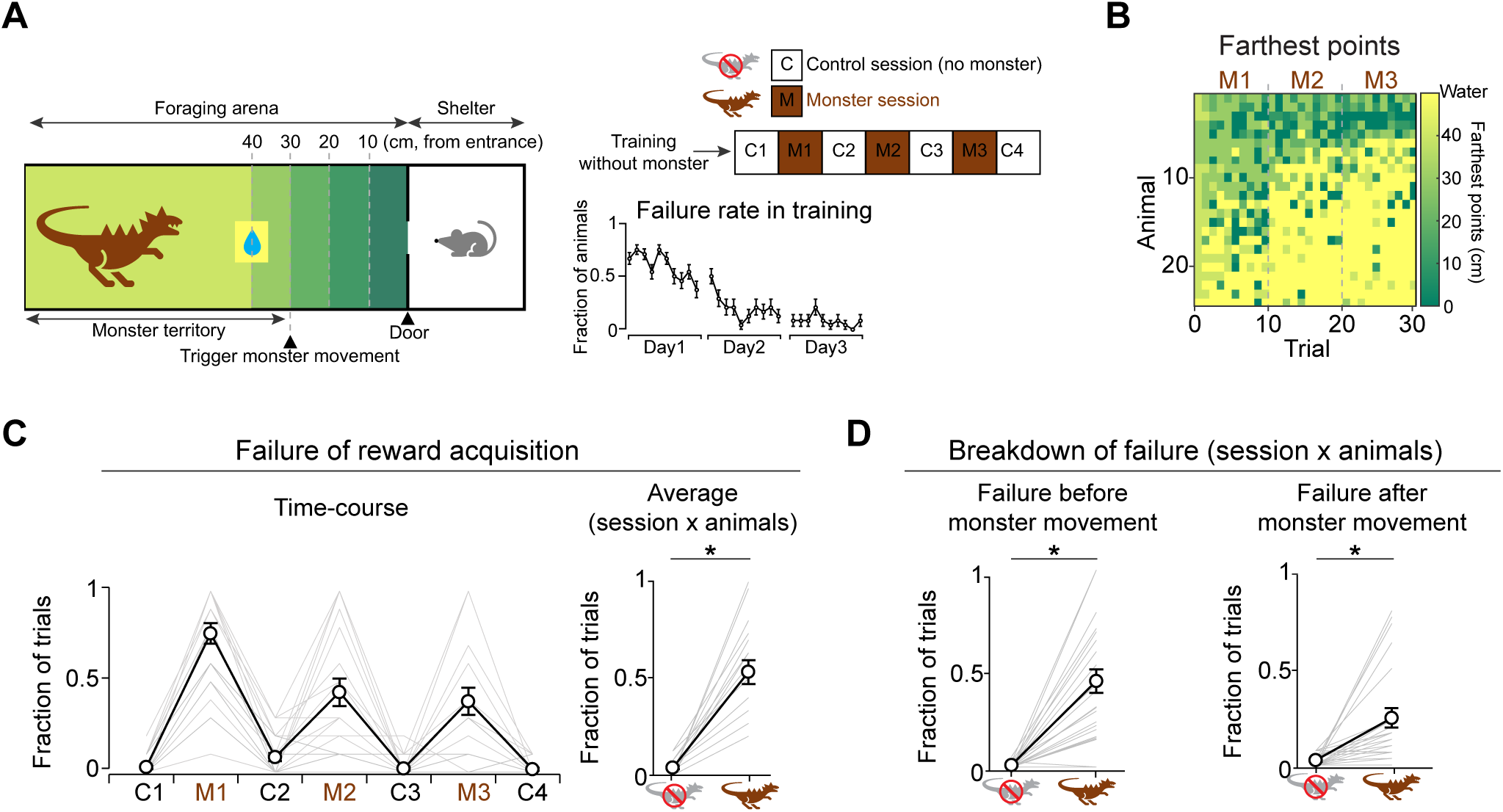
A “monster” paradigm where a mouse navigates a potential threat to acquire water reward. (**A)** Left, Monster apparatus. Monster movement was triggered when a mouse entered the “monster territory”. Right top, mice were trained without monster for 3 days, and then received 7 test sessions. Right bottom, the failure rate for reward acquisition gradually decreased across sessions during training (failure rate in day1, 0.6 ± 0.06; in day3, 0.08 ± 0.05, mean ± SEM, n=24 animals). (**B**) Heat map of the farthest points that each mouse reached per trial. The farthest points gradually decreased (i.e. return home earlier) within M1. Across sessions, gradually more mice succeeded. (**C**) Left, the time-course of the failure rate for reward acquisition. Right, the average failure rate was higher in the presence of the monster than in the control sessions (M1-M3 vs C1-C4, p=3.3×10^-8^, paired t-test). (**D**) The average rates of failure before entering the monster territory (failure before monster movement, p=2.7×10^-4^, paired t-test) and after entering the monster territory (failure after monster movement, p=1.0×10^-6^, paired t-test) were higher in the presence of the monster than in the control sessions.

We first examined dynamics of behaviors when mice foraged under a reward-threat conflict in the monster paradigm. Thirsty mice were first habituated in the shelter for 1 day, and then received 3 training sessions to harvest reward in the foraging arena without a monster (1 session/day, 10 trials/session). Animals quickly learned the task structure (Figure 1A, right bottom). After this training, animals then performed 7 sessions, alternating between sessions with and without a monster (Figure 1A, right top). The first session (C1) was always without a monster, and used as the baseline. In sessions with a moving monster, mice failed to harvest reward in a large fraction of trials with the failure rate gradually recovering over sessions (M1, M2, M3; Figure 1B and 1C, left). There are different failure types (Figures 1D and S1). In some trials, mice did not enter the foraging arena at all, although the frequency of this failure type is generally low and did not change across sessions. Almost all animals entered arena even in monster sessions (23/24 animals, Figure S1A). In many trials, mice advanced into the monster territory but often failed to acquire the water reward in the presence of a moving monster (“failure after monster movement”; Figure 1D, right). Additionally, mice sometimes did not enter the monster territory and returned to the shelter without triggering a monster movement (“failure before monster movement”; Figure 1D, left).

We next examined the time-course of behavioral changes (Figure 2). The very first time that a moving monster was presented, most mice (22/24 animals) went back to the shelter without harvesting reward (Figure 2A, left). In these trials, all mice advanced into the monster territory, which triggered the movement of the monster (Figure 2A, center) but then failed to make it to the reward location (Figure 2A, right). When we instead presented a motionless ’static’ monster, most mice successfully harvested a reward (Figure 2E, left), indicating that the movement of the monster was critical for their failures.

**Figure 2.**
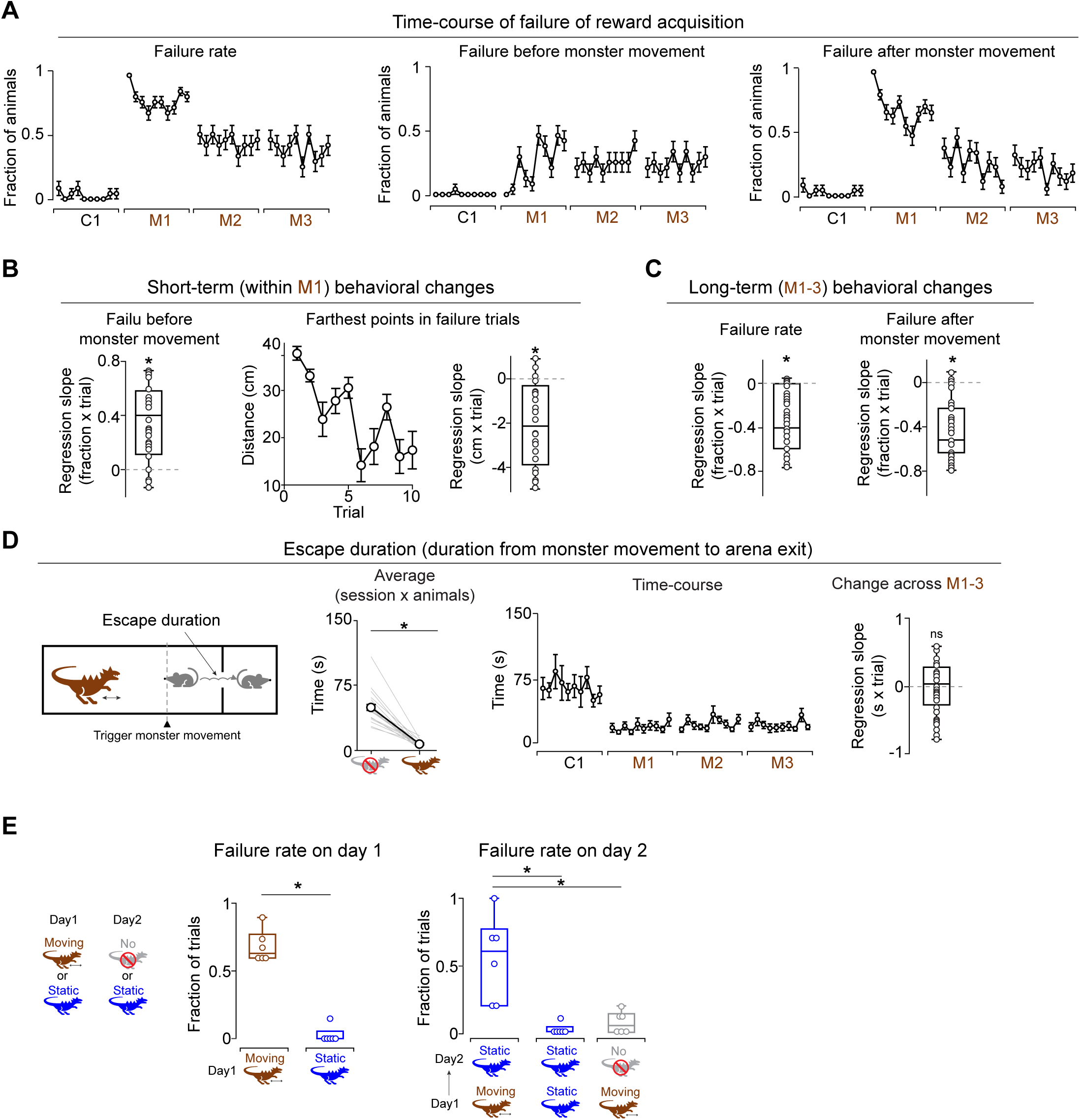
Dynamics of threat management behaviors in the monster paradigm. (**A**) Time-course of failures. Left, failure rate for reward acquisition in the first trial of M1 was significantly higher than in the first trial of C1 (χ^2^=25.08, p=5.5×10^-7^, chi-squared test). Center, rate of failure before monster movement. All mice succeeded in entering the monster territory in the first trial of M1. Right, rate of failure after monster movement in the first trial of M1 was significantly higher than in the first trial of C1 (χ^2^=25.08, p=5.5×10^-7^, chi-squared test). Error bars, SEM (binomial). (**B**) Left, regression coefficients of the rate of failure before monster movement as a function of trial number in M1 for each animal. The rate of failure before monster movement gradually increased across trials in M1 (p=6.1×10^-6^, t-test). Center, the farthest points in error trials gradually decreased across trials. Right, regression coefficients of the farthest points with trial number in M1 for each animal (p=1.9×10^-5^, t-test). (**C**) Regression coefficients of the failure rates for water acquisition (left, p=2.3×10^-7^, t-test) and the rate of failure after monster movement (right, p=4.9×10^-8^, t-test) with trial number in monster sessions (trials in M1, M2 and M3) for each animal. Failure rates gradually decreased across monster trials. (**D**) Escape duration is defined as the duration from the time when mouse entered the monster territory (i.e. monster movement) to the time when mouse exited the foraging arena. Second left, escape duration was shorter in the presence of the monster than in control sessions (p=3.6×10^-7^, paired t-test). Second right, time-course of the escape duration. Right, regression coefficients of escape duration with trial number in monster sessions. Escape duration was not significantly changed (p=0.87, t-test). (**E**) Left, failure rate was lower when a static monster was presented than when a moving monster was presented (p=2.5×10^-7^, t-test). Right, mice that had previously experienced a moving monster failed to acquire reward in the presence of a static monster on the next day (Moving→Static) more than mice who had previously experienced a static monster (Static→Static) (p=2.1×10^-3^, t-test). Mice that had experienced a moving monster failed less on the next day with no monster (Moving→No) than in the presence of static monster (Moving→Static) (p=4.5×10^-3^, t-test).

Further analysis showed that various additional aspects of behavior changed over trials. The failure of entering the monster territory slowly increased in the first session (Figure 2A, center and 2B, left). Similarly, the latency to enter the arena and to enter the monster territory gradually became longer, indicative of greater hesitation (Figure S1B and S1C). The farthest points that mice reached in the arena gradually became shorter (Figures 1B, M1 and 2B, center and right). These results suggest that the failure was initially triggered by the moving monster, but in later trials, mice predicted a potential threat and modulated their behavior accordingly even before triggering any monster movement in these trials.

The above observations suggest that in later trials, failures were increasingly caused by threat prediction rather than being directly triggered by a moving monster. We next examined in greater detail what caused this behavior. We reasoned that after experiencing a moving monster, sight of the monster was itself sufficient to increase hesitation in approach behavior. To test this idea, we probed the animal’s behavior using a static monster at different stages in the paradigm. We found that the static monster alone was not sufficient to evoke strong avoidance behavior in mice who had not yet experienced trials with the moving monster (Figure 2E, left). However, after mice were exposed to a moving monster, the presence of a static monster led to failures on most trials (Figure 2E, right, Moving→Static). If a static monster was presented first, mice successfully acquired reward in the next session when presented with a static monster (Figure 2E, right, Static→Static). Although mice showed markedly greater avoidance after encountering the moving monster, it is possible that experience might have evoked a general fear response that inhibited foraging ability, rather than failures resulting from a specific threat prediction attributed to the monster. To test this alternative hypothesis, we compared behaviors with or without a static monster after mice had already experienced the moving monster. We found that mice nearly always succeeded in capturing reward in sessions that did not include a monster, despite having experienced the moving monster in the previous session (Figure 2E, right, Moving→No). Together, these results indicate that mice learned to predict a potential threat in response to sight of a monster, even when it was motionless, after they had previously experienced it moving.

In parallel with the behavioral change within the first monster session (increasing avoidance before entering the monster territory in later trials), we also observed that the failure rate decreased slowly in a longer time scale (Figure 1B and 2A, left). Over multiple sessions, both overall failure of reward acquisition and failure after monster movement gradually decreased (Figure 2A, left and right, and 2C). Similarly, the latency to enter the arena and the monster territory gradually decreased (Figure S1D). However, other aspects of behavior suggest that the threat was not completely extinguished. We observed that mice quickly returned to the shelter after triggering a monster movement in monster sessions compared to sessions with no monster (“escape duration”; Figure 2D, left and second left). Interestingly, the escape duration remained short even after the success rate started to improve (Figure 2D, second right and right). Thus, mice continued to escape quickly from a moving monster at the similar level but did not suppress acquisition of reward. These results suggest that mice overcame a monster threat sufficiently to capture reward rather than extinguished the threat prediction associated with the monster altogether.

In summary, we observed a gradual development of threat management behaviors in our paradigm, transitioning across different components: threat avoidance, threat prediction, and overcoming threat. Importantly, mice never experienced a painful outcome, and dynamically adjust their avoidance behaviors of a potential threat, while maintaining quick return to the shelter.

### Intensity-coding and threat avoidance with dopamine in TS

Previous studies found that dopamine in TS plays a role in avoidance of a threatening stimulus (Akiti et al., 2021; Menegas et al., 2018). Activity of dopamine in TS did not reliably signal reward value but was instead monotonically modulated by intensity of external stimuli (Menegas et al., 2018; Menegas et al., 2017). While these studies suggested that different dopamine neurons send signals along two major axes: reward value and stimulus intensity, it was not clear whether intensity-coding is unique to dopamine in TS or shared with surrounding areas. Other studies have suggested that dopamine in both TS and the dorsolateral striatum (DLS) signals salience (Cox and Witten, 2019; Lerner et al., 2015; Matsumoto and Hikosaka, 2009; Menegas et al., 2017), and these two regions are often grouped together (Collins and Saunders, 2020). In order to examine the precise location of intensity-coding by dopamine in the striatum, we systematically mapped dopamine activity throughout the striatum along two axes: value and stimulus intensity.

Mice were injected with adeno-associated virus (AAV) to express the dopamine sensor, GRAB_DA2m_ (Sun et al., 2020) in the various striatal subareas (Figure 3). We mapped the dopamine activity patterns by recording the dopamine sensor signals in the striatum while head-fixed mice received various intensity of tone (50, 75, and 100 dB) and various amounts of water (1, 3, and 10 μl) in a pseudorandom order (Figure 3A). Overall, we observed a widespread excitation to water, while responses to tones varied in their sign across areas (Figure 3B). Dopamine activity was inhibited by tones in the anterior striatum but was activated in the posterior striatum (Figure 3B, bottom right). We quantified the activity modulation by reward amounts and tone intensity using linear regression (Figure 3C). The slopes of the correlations (beta coefficients) were compared across recording locations. Modulation by tone intensity was significantly correlated with anterior-posterior positions (Figure 3C, right), whereas modulation by water amounts did not show a significant trend but was slightly larger in the anterior regions (Figure 3C, left). Interestingly, water amount-coding and intensity-coding showed negative correlation with each other (Figure S2A). We then categorized activity patterns into 5 types: positive modulation with water amounts (value (+)), tone intensity (intensity) and both (value (+) & intensity), positive modulation with water and negative modulation with tone (value (+/–)), and others (Figure 3D). The spatial distribution of these response types indicated a clear separation between the anterior and posterior striatum. The anterior striatum (anterior to Bregma -0.5 mm) had value (+) and value (+/–) types, while the posterior striatum had intensity and value (+) & intensity types (Figure 3D, bottom). The boundary that separates these anterior and posterior regions did not correspond directly to the boundary based on cortical projection patterns (Hunnicutt et al., 2016) (Figures 3D-E and S2B). The observation of value (+) & intensity type at the boundary of DLS and TS (Bregma from -0.5 to -1.5 mm) is consistent with previous studies that observed excitation with both reward and high intensity stimuli in DLS or TS (Lerner et al., 2015; Menegas et al., 2017). While this boundary area was categorized as DLS in one study (Hunnicutt et al., 2016), other studies categorized it as TS because of unique developmental and biochemical characteristics different from more anterior DLS (Matsushima and Graybiel, 2020; Miyamoto et al., 2019; Valjent and Gangarossa, 2021). In monkeys, TS consists of the caudate tail and putamen tail (Amita et al., 2019; Kunimatsu et al., 2019). Since rodent DLS is anatomically similar to monkey putamen, our results, together with previous results, suggest that DLS/TS boundary area corresponds to the putamen tail in monkeys, and thus we tentatively call it tail of DLS or anterior TS (tDLS/aTS), although further verification would be needed to determine the exact boundaries because our recording method does not distinguish actual value (+) & intensity coding versus contamination of these two different signals at the boundary (Figure 3F). In this study, we will mainly focus on the posterior part of the intensity-coding area (posterior to Bregma -1.5 mm) as TS, in comparison with adjacent central parts of DLS (cDLS), anterior to Bregma -0.5 mm.

**Figure 3.**
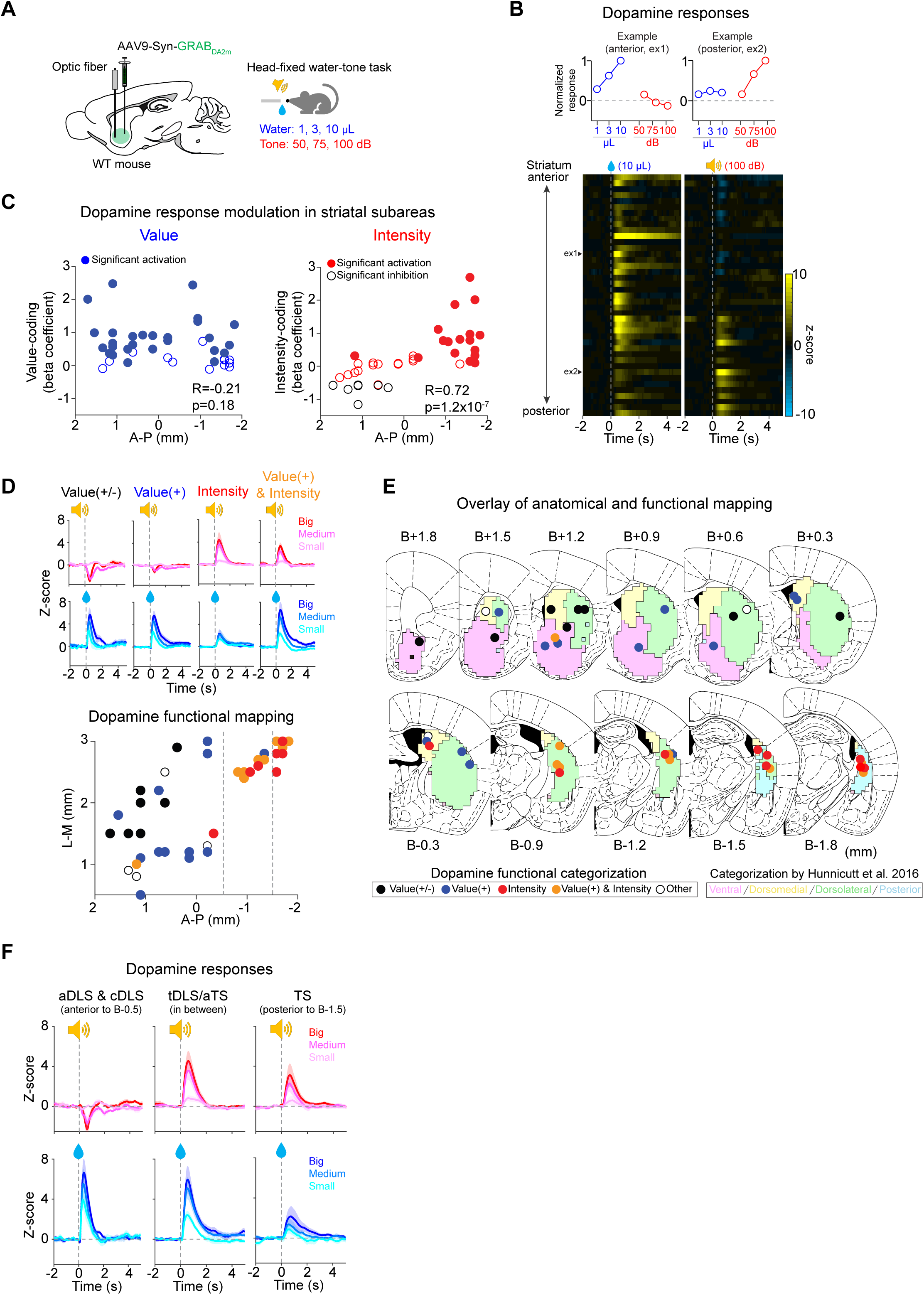
Value- and intensity-modulation of dopamine responses in the striatum. (**A**) Left, AAV9-Syn-GRAB_DA2m_ was injected, and an optic fiber was implanted in various areas of the striatum (43 animals, one recording site per animal). Right, during the fiber fluorometry recording, 3 different intensities of pure tone (50, 75, and 100 dB) and 3 different sizes of water (1, 3, and 10 μl) were pseudo-randomly presented to the head-fixed mice. (**B**) Top, dopamine responses (0-1s) to water and tone in example animals (left, anterior, ex1; right, posterior, ex2). Bottom, dopamine responses to water (left, 10 μl) and to tone (right, 100 dB) across different locations of the striatum. Example animals are indicated by arrowheads. (**C**) Regression coefficients of dopamine responses with water sizes (left) and tone intensities (right) were plotted along the anterior-posterior axis in the striatum. (**D**) Categorization of dopamine responses into value (+/–), value (+), intensity, and value & intensity types (mean ± SEM). Bottom, dopamine response types on an anatomical map of the striatum. (**E**) Dopamine response types were mapped onto coronal sections where the striatum was divided into 4 subareas using cortico-striatal projection patterns (Hunnicutt et al., 2016). (**F**) Dopamine responses to tone (top) and water (bottom) in anterior and central DLS (aDLS & cDLS, left), tail of DLS/anterior TS (tDLS/aTS, center), and TS (right) (mean ± SEM).

We next compared dopamine activity patterns in TS and cDLS in the monster paradigm (Figures 4 and S3). We first examined responses to water in no-monster sessions and responses to monster movement in monster sessions. Consistent with recording results in head-fixed mice, TS dopamine strongly responded to monster charge, while cDLS dopamine strongly responded to water reward (Figure 4A and 4B). Since cDLS dopamine showed slight excitation with some delay after monster charge (Figure 4A, bottom right), we next performed a regression analysis to fit multiple kernels aligned to water delivery or monster movement onset with dopamine activity in each animal to construct activity model (see Methods). This method can separate responses triggered by temporally overlapping events. We found that cDLS dopamine responses were mainly explained by water delivery, while TS dopamine responses were mainly explained by monster charge (Figures 4C and S3), although TS dopamine models were more variable across individuals (see below).

**Figure 4.**
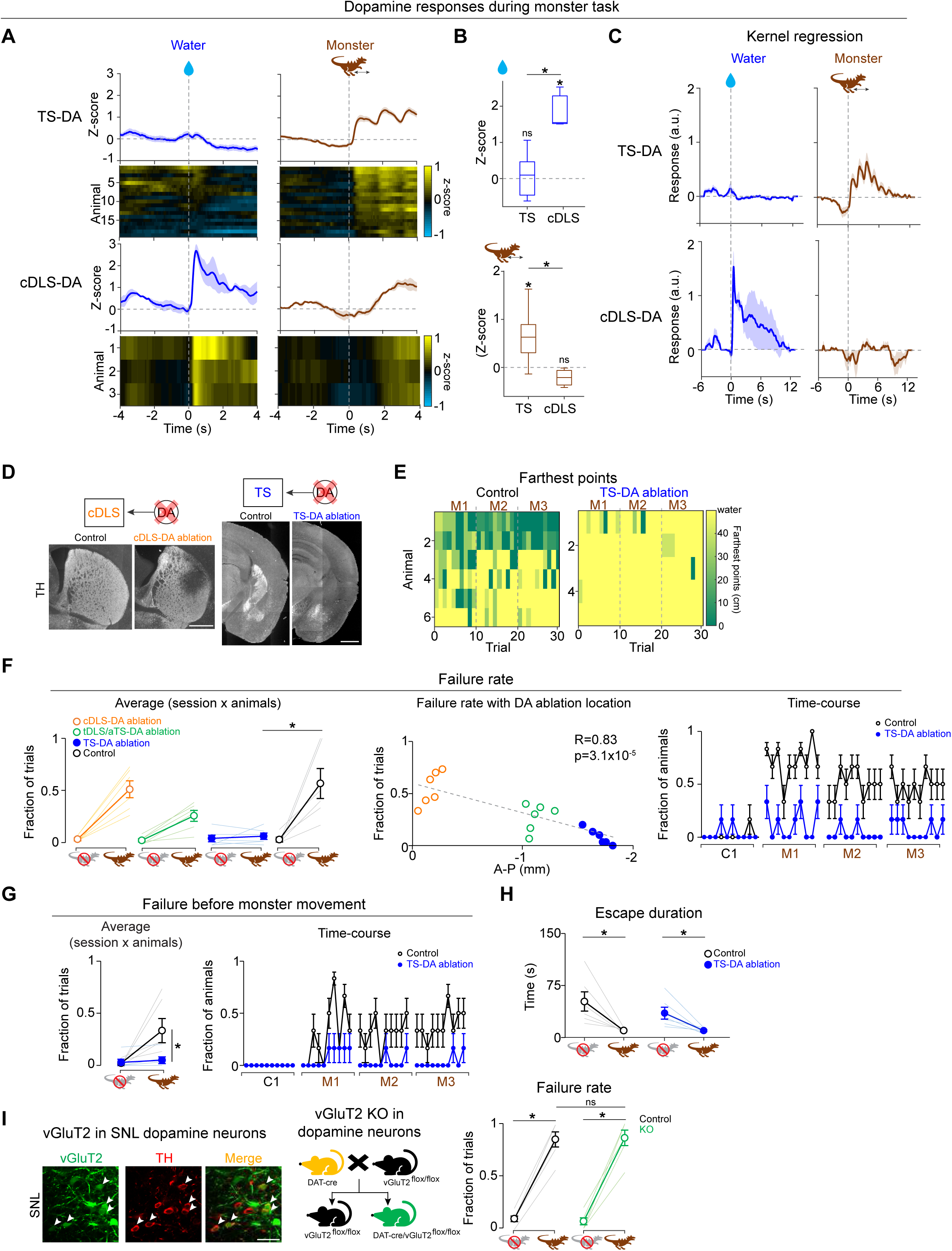
TS dopamine plays a role in threat avoidance and prediction. (**A**) Dopamine sensor signals in TS (n=19 animals, top) and cDLS (n=3 animals, bottom) during the monster task. Left, dopamine responses to water in no monster baseline session. Right, dopamine responses to monster movement in monster sessions M1-3 (right) (mean ± SEM). (**B**) Average dopamine responses (0-1 s) to water (top) and monster movement (bottom). Dopamine sensor signals in cDLS showed significant activation with water (p=8.0×10^-3^, t-test) but not in TS (p=0.58, t-test) and the water responses were significantly higher in cDLS than in TS (p=6.2×10^-7^, t-test). Dopamine sensor signals showed significant activation at monster movement in TS (p=3.0×10^-5^, t-test) but not in cDLS (p=0.56, t-test), and the monster responses were significantly higher in TS than in cDLS (p=0.021, t-test). (**C**) Average responses of kernel regression models (mean ± SEM). (**D**) Ablation of dopamine neurons with 6-OHDA. White, anti-tyrosine hydroxylase (TH). Bar, 1 mm. (**E**) Heat map of the farthest points that control animals (left) and animals with ablation of TS-projecting dopamine neurons (TS-DA ablation, right) reached in each trial. (**F**) Failure rate for reward acquisition. Left, the average failure rate in cDLS-, tDLS/aTS-, and TS-DA ablation mice, and control mice (n=6, each). Failure rates were modulated by anterior-posterior location of dopamine ablation (F(3,23)=6.29, p=4.0×10^-3^, ANOVA). In the presence of the monster, failure rate was significantly higher than in control sessions in control mice (p=0.016, paired t-test) but not in TS-DA ablation mice (p=0.45, paired t-test), and failure rate in monster sessions was higher in control mice than in TS-DA ablation mice (p=0.022, t-test). Center, failure rate in monster sessions and dopamine ablation locations in the striatum. Failure rate was higher in mice with dopamine ablation in more posterior parts of the striatum (R=0.83, p=3.1×10^-5^, Pearson’s correlation coefficients). Right, time-course of failure rate in control and TS-DA ablation mice. TS-DA ablation mice succeeded in reward acquisition from the first trial of M1 (first trial in M1 vs. first trial in C1, χ^2^=6, p=0.014, chi-squared test). Error bars, SEM (binomial). (**G**) Left, average rate of failure before monster movement was higher in monster sessions than in control sessions in control mice (black, p=0.048, paired t-test) but not in TS-DA ablation mice (blue, p=0.088, paired t-test), and failure rate in monster sessions was higher in control mice than in TS-DA ablation mice (p=0.047, t-test). Right, time-course of rate of failure before monster movement in control and TS-DA ablation mice. Error bars, SEM (binomial). (**H**) Escape duration was significantly shorter in monster sessions than in control sessions with both control mice (black, p=2.7×10^-3^, paired t-test) and ablation mice (blue, p=5.8×10^-3^, paired t-test). (**I**) Behavioral analysis using dopamine neuron-specific conditional knockout (KO) of the vGluT2 gene. Left, representative image of vGluT2 (green) and tyrosine hydroxylase (TH, red) positive neurons in SNL. Co-localization is indicated by white arrow. Bar, 20 μm. Right, average failure rate for reward acquisition in control and KO mice. Both control (p=3.2×10^-5^, paired t-test) and KO (p=9.3×10^-5^, paired t-test) failed more in monster sessions than in control sessions, and failure rates in monster sessions in control mice and KO mice were similar (p=0.96, t-test).

The strong excitation by monster movement suggested a role of TS dopamine in avoidance. We next compared the function of dopamine in cDLS and TS by ablation of dopamine neurons that project to different striatal areas using targeted injection of 6-hydroxydopamine (6-OHDA) (Figure 4D-H). We found that ablation of TS-projecting dopamine neurons reduced the failure of reward acquisition, while ablation of cDLS-projecting dopamine neurons did not affect it (Figure 4F, left). This ablation effect (suppression of failure rate) was strongly correlated with ablation locations along the anterior-posterior axis (Figure 4F, center). Notably, mice with TS-projecting dopamine neuron ablation successfully harvested reward from the first encounter with the moving monster (Figure 4E-G). On the other hand, the escape duration did not change by ablation (Figure 4H); both control (Figure 4H, black) and ablation (Figure 4H, blue) mice exited the arena more quickly in the presence of a monster than in control sessions. These results suggest that the ablation mice recognized the moving monster, yet they advanced to the reward location, harvested the reward, and then quickly escaped.

Although these results suggest that dopamine in TS is important for avoidance in a monster paradigm, ablation studies do not distinguish roles of different neurotransmitters. A previous study found that dopamine neurons in the substantia nigra pars lateralis (SNL), where the cell bodies of TS-projecting dopamine neurons are mainly localized (Menegas et al., 2015), express vesicular glutamate transporter 2 (vGluT2) (Poulin et al., 2018; Yamaguchi et al., 2013) (Figure 4I, left), suggesting co-release of glutamate and dopamine in TS. We next tested whether glutamate release from dopamine neurons is important for threat avoidance. We genetically removed vGluT2 expression in dopamine neurons using dopamine transporter (DAT)- cre/vGluT2^lox/lox^ mice (Figure 4I, center). Removal of vGluT2 in dopamine neurons did not affect the failure rate (Figure 4I, right). Thus, glutamate release from dopamine neurons is dispensable for threat avoidance.

Because we found that dopamine activity in TS is uniquely modulated by stimulus intensity, and that TS dopamine facilitates avoidance of a potential threat, in the following, we will focus on TS to examine the mechanism of threat management in more detail.

### Modulation of avoidance behaviors by dopamine in TS according to threat size

We found that TS dopamine plays a role in avoidance of a potential threat (Figure 4) and uniquely signals stimulus intensity (Figure 3). How does TS dopamine function in threat avoidance? What is the relationship between stimulus intensity and threat avoidance? We previously proposed that TS dopamine signals physical salience of a stimulus as an initial estimate of threat level (Akiti et al., 2021), although the idea had not been tested experimentally. To address this, we next examined the relationship between physical salience of a threatening stimulus, TS dopamine activity, and avoidance behaviors (Figure 5A-D). We first examined whether the size of the monster affects animals’ avoidance behaviors. Different groups of mice were tested in the monster paradigm with a big (18 cm, same size as Figure 1), medium (10 cm) or small (3 cm) monster (Figure 5A). We found that the failure rate for reward acquisition was affected by monster size; mice failed significantly more with the big monster than with the small monster (Figure 5A, bottom, left). This was not because mice did not notice the small monster. Regardless of whether the monster was big or small, the escape duration was similarly shorter in monster sessions (Figure 5A, bottom, right), indicating that mice noticed the monster and changed their behaviors in both cases. However, in the case of the small monster, mice did not suppress water acquisition; they quickly ran away only after drinking water. Thus, mice modulated their behavior differently based on the size the monster.

**Figure 5.**
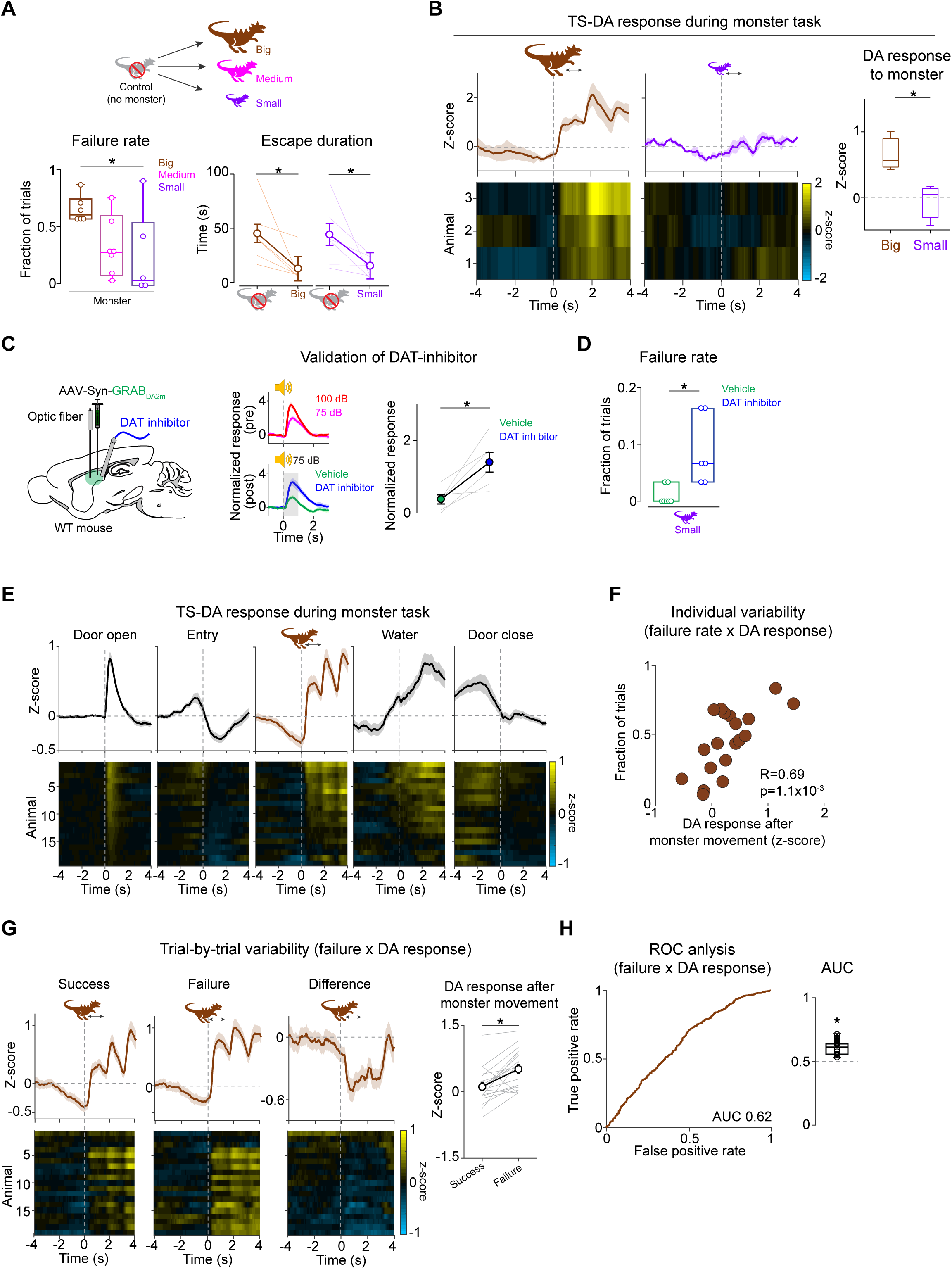
TS dopamine signals are modulated by physical salience of a potential threat and facilitate avoidance. (**A**) Top, different groups of mice performed the monster paradigm with different sizes of monster (n=6 animals, each). Bottom left, the average failure rate for reward acquisition in the monster session. The monster’s size modulated the failure rate (F(2,17)=5.55, p=0.040, ANOVA). Failure rate with the small monster was significantly lower than with the big monster (p=0.030, t-test). Bottom right, the average escape duration was significantly shorter in the monster session than in the control session in both big and small monster groups (big, p=0.032; small, p=3.4×10^-3^, paired t-test). (**B**) Left, dopamine sensor signals at movement of a big monster (left, n=3 animals) or a small monster (center, n=3 animals) (mean ± SEM). Right, average dopamine responses at movement of a big or small monster (0-1 s). Dopamine responses to a big monster were significantly higher than responses to a small monster (p=0.042, t-test). (**C**) Validation of a dopamine transporter (DAT) inhibitor, GBR12909. Left, AAV9-Syn-GRAB_DA2m_ was injected into TS. Dopamine sensor signals were recorded while head-fixed mice were presented with a complex tone. Center, dopamine responses to tone before (top, “pre”) and after (bottom, “post”) injection of DAT inhibitor or vehicle. Dopamine responses were normalized by the responses to 75 dB of tone in the pre-injection session. Right, average dopamine responses to tone (0-1s). Dopamine sensor signals were significantly higher when DAT inhibitor was infused in the TS than vehicle (p=0.011, n=6 sessions with 3 animals, paired t-test). (**D**) DAT inhibitor (n=6 animals) or vehicle (n=6 animals) was bilaterally injected into TS, and mice were tested in the monster paradigm with a small monster. Failure rate for reward acquisition was significantly higher with DAT inhibitor in TS than with vehicle (p=0.014, t-test). (**E-H**) Dopamine sensor signals in TS during the monster task (n=19 animals). (**E**) Top, average dopamine responses across animals (mean ± SEM) locked to different task epochs. Bottom, dopamine responses in each animal. (**F**) Average TS dopamine responses to monster were positively correlated with individual variability of failure rate for reward acquisition (R=0.69, p=1.1×10^-3^, Pearson’s correlation coefficient). (**G**) Dopamine responses at the monster movement in success trials (left), failure trials (second left), and the difference of those (second right) (mean ± SEM). Right, dopamine responses at monster movement were significantly higher in failure than in success trials (p=1.1×10^-4^, paired t-test). (**H**) Left, receiver operating characteristic (ROC) curve evaluating the discriminability of dopamine responses at monster movement between success and failure trials (AUC=0.62). Right, AUC for each animal. (0.60 ± 0.05, mean ± STD; AUC vs 0.5, p=5.8×10^-5^, t-test).

Next, we examined whether TS dopamine activity is similarly varied with size of monster. TS dopamine activity was recorded while different groups of mice performed in the monster paradigm with either a big or small monster. We found that TS dopamine was strongly activated by movement of the big monster, but not the small monster (Figure 5B). Thus, we observed correlation between threat size and dopamine activity, as well as between threat size and avoidance behaviors.

We next examined causality. We showed that ablation of TS-projecting dopamine neurons suppressed the failure of reward acquisition in the presence of a big monster (Figure 4F). We next performed the opposite manipulation: does artificial inflation of dopamine signals exaggerate effects of threat size? In order to boost dopamine signaling in a physiologically relevant range, we utilized the dopamine transporter (DAT) inhibitor (GBR12909), which inhibits dopamine reuptake from the extra-cellular space, causing an increase of lingering dopamine. We first examined effects of DAT inhibitor on dopamine signaling in TS (Figure 5C), since DAT expression is lower in TS (Gangarossa et al., 2013; Miyamoto et al., 2019). Dopamine activity was examined while head-fixed mice were exposed to different intensity of tones (Figure 5C, center top). After confirming intensity-modulation in dopamine signals, either DAT inhibitor or vehicle was infused into TS. We confirmed that dopamine sensor signals to tones were increased by DAT inhibitor (Figure 5C, center bottom and right), although our method likely underestimates the effect size due to baseline increase in dopamine levels. We next examined the effect of boosting dopamine signaling on avoidance behaviors. Different groups of cannulated mice were bilaterally infused with either DAT inhibitor or vehicle, and were then tested in the monster paradigm with a small monster. While most mice with vehicle infusion successfully acquired reward with a small monster, DAT infusion significantly increased the failure rate (Figure 5D). This result demonstrates a causal role of dopamine signaling in avoidance behavior that depends on the size of a threatening stimulus. Taken together with TS dopamine activity patterns (monotonic modulation with intensity of various stimuli), these results imply that TS dopamine signals an estimate of a potential threat according to the physical salience of a stimulus.

Since artificial manipulation of dopamine signaling affected avoidance behaviors, we next examined natural individual variability in dopamine activity and behaviors. There is a wide range of individual variability in avoidance behaviors in our monster paradigm (Figure 1B-D).

Dopamine responses to monster movement were also variable (Figure 5E, center). Interestingly, across mice, the average dopamine responses to the moving monster were positively correlated with the average failure rate for reward acquisition (Figure 5F). This observation further supports the idea that TS dopamine signals causally regulate avoidance behaviors.

How does dopamine in TS facilitate avoidance by signaling physical salience? It has been shown that canonical dopamine neurons send reward prediction error as evaluation signals to update reward prediction (Bayer and Glimcher, 2005; Cohen et al., 2012; Schultz et al., 1997). Do the physical salience signals in TS dopamine also function as evaluation signals to reinforce threat prediction and avoidance in a similar manner as canonical dopamine does for reward? A previous study found that optogenetic activation of TS-projecting dopamine neurons also showed reinforcing effects on avoidance of a novel object, suggesting development of threat prediction (Menegas et al., 2018). We next examined whether ablation of TS-projecting dopamine neurons affected the behavior that would depend explicitly on threat prediction, focusing on whether mice returned to the shelter without triggering monster movement (“Failure before monster movement”). Control mice showed a gradual increase of failure of entering the monster territory in the first session (Figure 4G, right, black). Conversely, ablation mice did not show this increase (Figure 4G, right, blue), indicating that TS dopamine is important for threat prediction based on the sight of the monster.

However, learning threat predictions might not explain all the behavioral effects of dopamine since the ablation effect manifested even in the very first trial (Figure 4E-F). In order to understand effects of dopamine on the current trial, we next examined trial-to-trial correlation between natural variability in TS dopamine responses and failure. We found that responses to monster movement in failure trials were significantly greater than in success trials (Figure 5G). Dopamine responses to monster movement were predictive of failure in the current trial, as characterized using the receiver-operating characteristics (ROC) analysis (Figure 5H). Whether these trial-based effects are directly caused by acute trigger of avoidance, or indirectly by moment-by-moment updates of threat prediction is an important question, but one that is difficult to tease apart.

Our results demonstrated that TS dopamine signals physical salience of a stimulus, leading to a facilitation of avoidance and prediction of a potential threat. The behavioral variability across animals as well as on a trial-by-trial basis were partially explained by dopamine responses triggered by the monster.

### Direct pathway neurons in TS are important for avoidance

The above results indicate that dopamine in TS is important for threat avoidance in the current trial and threat prediction in later trials. What is the function of the downstream neurons in TS? Similarly to other striatal areas, projection neurons in TS are categorized into two cell types depending on expression of dopamine receptor type 1 (D1) or type 2 (D2) (Miyamoto et al., 2019; Valjent and Gangarossa, 2021). We used Tachykinin precursor 1 (Tac1)-cre mice and adenosine 2A receptor (Adora2A)-cre mice to manipulate D1- and D2-expressing projection neurons, respectively (’dMSN’ and ’iMSN’ hereafter), because D1 and D2 receptors are also expressed in local neurons in the striatum (Alcantara et al., 2003; Aosaki et al., 1998; Pisani et al., 2000). Previous studies found that dopamine strengthens the cortical synapses onto dMSNs (Iino et al., 2020; Shen et al., 2008; Yagishita et al., 2014) as well as activates dMSNs on a sub-second timescale (Lahiri and Bevan, 2020). These findings suggest the possibility that dMSNs in TS play a role in threat avoidance and threat prediction under reward-threat conflict, similar to dopamine in TS.

Following the completion of 3 sessions of training without the monster, we injected AAV to express diphtheria toxin subunit A (dtA) (Wu et al., 2014) bilaterally into TS in Tac1-cre/tdTom mice, to specifically ablate dMSNs in TS (Figures 6 and S4). After 14 days, behaviors were tested in the monster paradigm. Similar to ablating TS-projecting dopamine neurons, ablation of dMSNs in TS decreased the failure rate for reward acquisition (Figure 6B-C). This decrease in failure rate was significantly correlated with ablation location, determined by loss of tdTomato signals, along the anterior-posterior axis (Figure 6C, center); dMSNs in the posterior striatum (TS) were specifically important for avoidance, but not in more anterior cDLS. The difference between control and ablation animals was mainly explained by failure of reward acquisition after monster movement (Figure S4B). In addition, ablation animals exhibited a slight decrease in the average failure rate of entering the monster territory (Figure S4A), indicating that dMSNs are also important for threat prediction.

**Figure 6.**
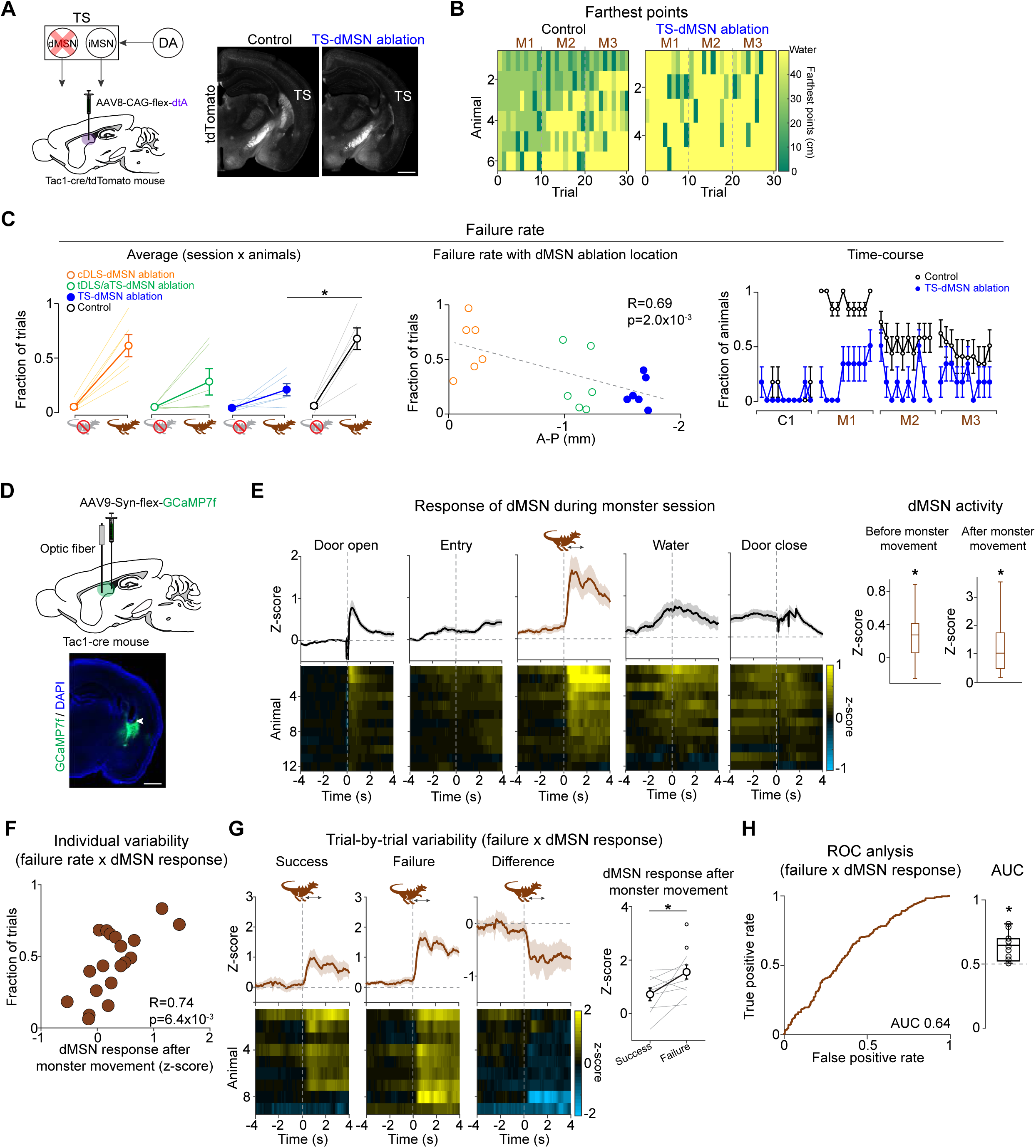
Direct pathway neurons in TS play a role in threat avoidance and prediction. (**A**) Right, AAV8-CAG-flex-diphtheria toxin subunit A (dtA) was bilaterally injected into the posterior striatum in Tac1-cre/Ai14 mice to specifically ablate dMSNs. Control mice received bilateral injection of saline into the TS. Left, representative histological images from control and ablation animals. White, tdTomato. Bar, 1 mm. (**B**) Farthest points that control (left) and TS-dMSN ablation (right) mice reached in each trial. (**C**) Left, the average failure rate for reward acquisition in cDLS-, tDLS/aTS-, and TS- dMSN ablation, and control mice (n=6 animals, each). The failure rate was significantly higher in monster sessions than in control sessions in the control mice (p=9.6×10^-3^, paired t-test) but not in TS-dMSN ablation mice (p=0.076, paired t-test). The failure rate was modulated by the anterior-posterior locations of dMSN ablation (p=0.014, F(3,23)=4.50, ANOVA). The failure rates in the monster sessions were significantly higher in the control mice than in TS-dMSN ablation mice (p=6.1×10^-4^, t-test). Center, mice with ablation of dMSNs in more posterior striatum showed lower failure rates in the presence of the monster (R=0.69, p=2.0×10^-3^, Pearson’s correlation coefficients). Right, the time-course of the failure rate in control and TS-dMSN ablation mice. Error bars, SEM (binomial). (**D-H**) Recording of population activity of dMSNs in TS during the monster task (n=12 animals). (**D**) Top, AAV9-Syn-flex-GcaMP7f was injected unilaterally into TS of Tac1-cre mice. Bottom, location of an optic fiber in an example animal. Arrowhead, the tip of a fiber. Green, GCaMP7f. Blue, DAPI. Bar, 1 mm. (**E**) the average dMSN activity across animals (mean ± SEM). Right, average responses (0-1s). dMSNs were activated before (p=0.012, t-test) and after the monster movement (p=1.3×10^-3^, t-test). (**F**) Average TS-dMSN responses to monster movement were positively correlated with individual variability of the failure rates for reward acquisition (R=0.74, p=6.4×10^-3^, Pearson’s correlation coefficient). (**G**) dMSN activity at monster movement in success trials (left) and failure trials (second left), and the difference of those (second right) (mean ± SEM). Right, dMSN responses to monster movement were higher in failure trials than that in success trials (p=0.033, paired t-test). (**H**) Left, ROC curve evaluating the discriminability of the dMSN responses at monster movement between success and failure trials (AUC=0.64). Right, AUC for each animal (0.63 ± 0.10, mean ± STD; AUC vs 0.5, p=5.7×10^-3^, t-test).

We next examined the population activity of dMSNs in the monster task using Ca^2+^ indicator GCaMP7f (Dana et al., 2019) (Figures 6D-H, S4C). In the baseline session before the first encounter with the monster, dMSNs were excited at door opening and closing (Figure S4C). After introduction of the monster, dMSNs were strongly activated after monster movement (Figure 6E). Similar to dopamine neurons, dMSN responses to monster movement were variable across animals (Figure 6E, center and right), and the average dMSN responses were positively correlated with individual variability of the failure rate for reward acquisition (Figure 6F). Further, dMSN responses to monster movement were correlated with failure on a trial-by-trial basis (Figure 6G), and dMSN responses to monster movement were predictive of failure in the current trial (Figure 6H). Thus, dMSNs in TS play a role in threat avoidance and threat prediction, similar to dopamine in TS, consistent with the previous findings that dopamine activates and strengthens dMSN signaling through D1 receptors.

### Indirect pathway neurons in TS are important for overcoming threat

While the facilitatory role of dMSNs in the striatum on output signaling is widely accepted (Cruz et al., 2020; Durieux et al., 2012; Kravitz et al., 2010; Kravitz et al., 2012; Natsubori et al., 2017; Tecuapetla et al., 2016; Vicente et al., 2016), the function of iMSNs has long been under a heated debate (Cruz et al., 2020; Lee and Sabatini, 2021; Natsubori et al., 2017; Soares-Cunha et al., 2016a; Tecuapetla et al., 2016). While biochemical and anatomical properties point to opposing roles of dMSNs and iMSNs (Albin et al., 1989; DeLong, 1990; Gerfen and Surmeier, 2011), similar activity patterns observed in some studies encouraged arguments against it (Cui et al., 2013; Natsubori et al., 2017; Parker et al., 2018; Soares-Cunha et al., 2016b). Further, anatomical complexity of the basal ganglia suggests that the role of these neurons may not be symmetrical, even if they work in opposition (Courtney, 2021; Kita, 2007). Because appropriate threat management depends on multitudes of behavioral controls as described above, which aspects of behavior are controlled by iMSNs, and how, remains unclear.

We, therefore, next examined the function of iMSNs in the monster paradigm (Figure 7 and S5). After ablation of iMSNs in TS, mice showed a similar level of failure at the beginning as control mice (Figure 7B-C, M1), suggesting that iMSNs were not necessary for threat avoidance in this task. Further, iMSN ablation mice learned threat prediction normally over trials, indicated by the gradual increase of failure of entering the monster territory (Figure S5A). However, in stark contrast to control mice, iMSN ablation mice did not improve at obtaining reward over days (Figure 7C). While control mice showed a correlation between failure rate and trial number, iMSN ablation mice did not show such a correlation (Figure 7D, left). Lumping all the monster sessions, the failure rate was only slightly higher in iMSN ablation mice (Figure 7C, left), yet there was a large difference in the extent to which the failure rate decreased over trials, indicating that the ablation impaired the mice’s ability to overcome the potential threat. Importantly, the difference in the failure rate between ablation and control mice was mainly due to the increased failure *after* monster movement. While the failure after monster movement gradually decreased in controls, iMSN ablation mice did not show such improvement (Figure 7D, right), indicating that iMSN ablation mice did not overcome threat induced by monster movement.

**Figure 7.**
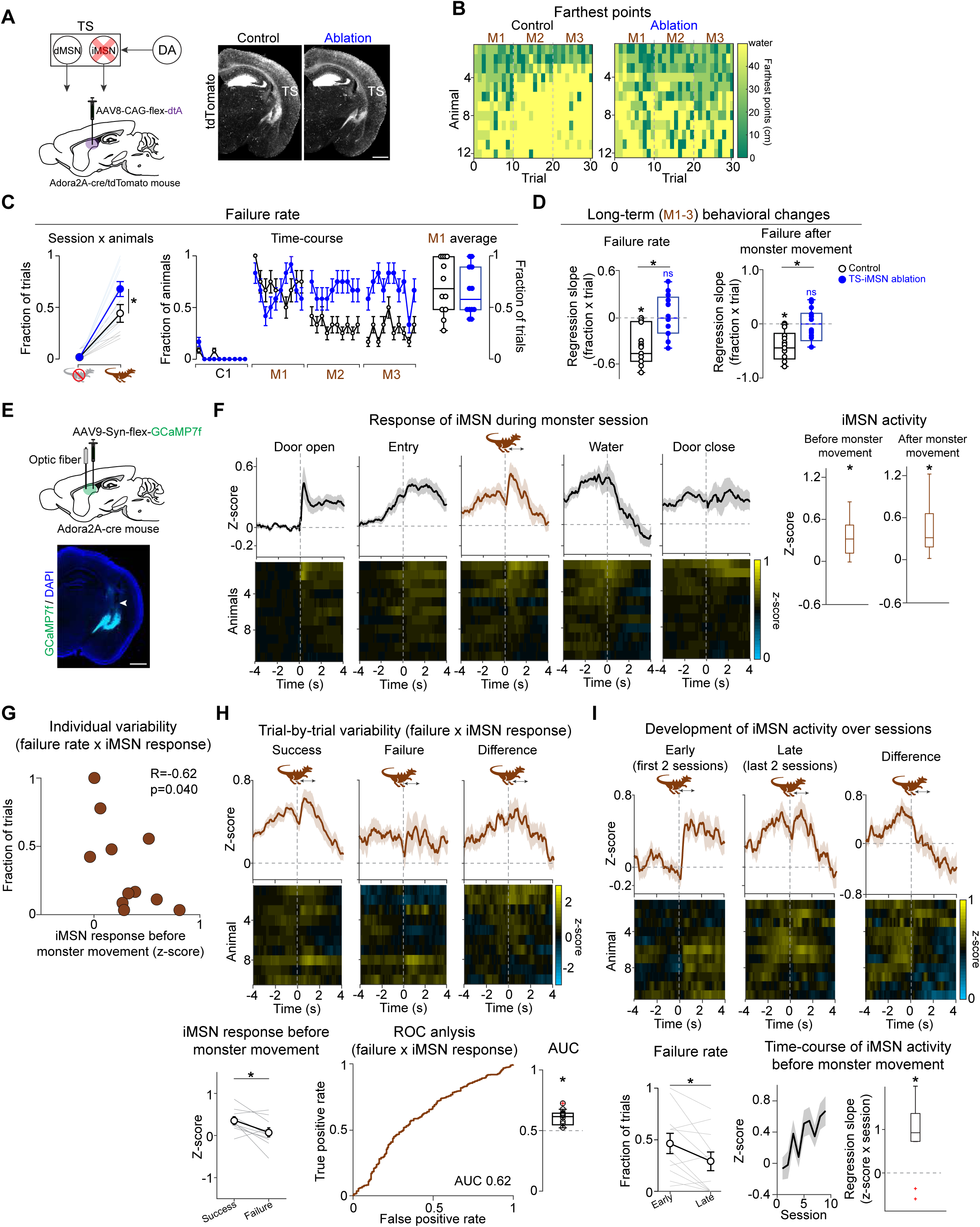
Indirect pathway neurons in TS play a role in overcoming a threat. (**A**) Left, AAV8-CAG-flex-dtA was bilaterally injected into the TS of Adora2a-cre/Ai14 mice (n=12 animals) to specifically ablate iMSNs. Control mice received bilateral injection of saline into the TS (n=12 animals). Right, representative histological images from control and ablation animals. White, tdTomato. Bar, 1 mm. (**B**) Heat map of the farthest points that control (left) and ablation (right) mice reached in each trial. (**C**) Left, the average failure rate for reward acquisition. Both control (p=5.8×10^-4^, paired t-test) and ablation (p=3.2×10^-6^, paired t-test) mice failed more in the presence of the monster than in control sessions. Failure rates were higher in ablation mice than control mice (p=0.048, t-test). Right, the time-course of the failure rate. Error bars, SEM (binomial). (**D**) The failure rate gradually decreased in control mice, but not in TS-iMSN ablation mice (left, regression coefficient of the failure rate with trials, control, p=6.4×10^-4^, ablation, p=0.61; control vs. ablation, p=4.4×10^-4^, t-test). The rate of failure after monster movement gradually decreased in control mice but not in ablation mice (right, regression coefficient of the rate of failure after monster movement with trials, control, p=1.6×10^-4^, ablation, p=0.63; control vs. ablation, p=2.3×10^-3^, t-test). (**E-I)** Recording of population activity of iMSNs in TS during the monster task (n=11 animals). (**E**) Top, AAV9-Syn-flex-GCaMP7f was unilaterally injected into TS of Adora2a-cre mice. Bottom, location of the optic fiber in an example animal. Arrowhead, the tip of the fiber. Green, GCaMP7f. Blue, DAPI. Bar, 1 mm. (**F**) Top, the average iMSN activity across animals (mean ± SEM). Bottom, iMSN activity in each animal. Right, average iMSN responses 0-1s before and after monster movement. iMSNs were activated before (p=2.9×10^-3^, t-test) and after the monster movement (p=1.6×10^-3^, t-test). (**G**) The average iMSN activity before monster movement was negatively correlated with individual variability of the failure rates for reward acquisition (left, R=-0.62, p=0.040, Pearson’s correlation coefficient). (**H**) Top, iMSN activity at the monster movement in success trials (left) and failure trials (center), and the difference of those (right) (mean ± SEM). Bottom, iMSN activity before monster movement was significantly higher in success than failure trials (left, paired t-test, p=0.024). ROC curve evaluating the discriminability of iMSN activity before monster movement between success and failure trials (AUC=0.62). Right, AUC for each animal (0.61 ± 0.08, mean ± STD; AUC vs 0.5, p=1.6×10^-3^, t-test). (**I**) Top, iMSN activity at the monster movement in early (left, session 1-2) and late (center, session 9-10) phases, and the difference of those (right) (mean ± SEM). Bottom, the failure rate was significantly lower in the late sessions than in the early sessions (left, p=0.024, paired t-test). Activity of iMSNs before monster movement increased over sessions (center, linear regression of average iMSN activity before monster movement with the session number, coefficient beta=0.088, p=2.5×10^-3^; right, beta coefficient of each animal, p=6.8×10^-3^, t-test).

The function of iMSNs in overcoming threat was different from function of dMSNs in threat avoidance and prediction in two fundamental ways. First, the directions of behavioral changes were opposite; ablation of dMSNs suppressed avoidance (increased successful reward acquisition), while ablation of iMSNs promoted avoidance (increased failure). Second, the effect of ablations manifested at different stages of threat management: initial threat learning versus later overcoming or suppressing of the threat. These results thus demonstrate fundamental differences in the roles that dMSNs and iMSNs in TS play in our paradigm.

We next examined activity patterns of iMSNs (Figure 7E-I). At a glance, activity patterns of iMSNs were similar to that of dMSNs. Before mice ever encountered the monster, iMSNs showed slight excitation with door opening and closing (Figure S5B). In monster sessions, iMSNs were activated after monster movement (Figure 7F). Notably, however, activation of iMSNs started even before the monster movement. The iMSN activation started right after the door opening, continued until onset of monster movement, and immediately dropped after water acquisition (Figure 7F). Such ramping activity was not observed in the baseline session in which no monster was present (Figure S5B, center), indicating that the excitation before monster movement was induced by the presence of a monster. We next examined correlation of iMSN activity and behavioral outcomes. In contrast to dopamine neurons and dMSNs, individual variability of iMSN responses before monster movement was negatively correlated with the failure rate for reward acquisition (Figures 7G); more iMSN activity was correlated with more success of the animal. Further, the activity before and after monster movement was correlated with success on a trial-to-trial basis (Figure 7H and S5C), and predictive of success (Figure 7H bottom right and S5D); iMSN activity before monster movement predicted upcoming success in water acquisition, suggesting the possibility that iMSNs proactively suppressed threat avoidance, i.e. the failure of obtaining reward.

Finally, consistent with the ablation results which showed an important role of iMSNs in overcoming threat in later trials, activity of iMSNs developed over trials around the time mice improved at obtaining reward (Figure 7I, bottom, left). Specifically, iMSN activity both at and before monster movement gradually increased over trials (Figure 7I and S5E). Together, these results indicate that dMSNs and iMSNs exert opposing functions in threat management – threat avoidance and overcoming, respectively – which are acquired at different stages over the course of learning.

## DISCUSSION

Although simple threat-triggered responses such as freezing and escape have long been studied intensively in neuroscience, how animals respond to, learn from, and eventually overcome a potential threat (“threat management”) is understudied. Threat management requires estimation of potential threats without actually experiencing ultimate outcomes, and flexible action-selection according to the threat estimates together with other factors such as rewarding opportunities.

In this study, we found that dopamine and the direct and indirect pathway neurons (dMSN and iMSN) in TS play critical roles at different stages of threat management under threat-reward conflict. Dopamine and dMSNs facilitate threat avoidance and prediction, while iMSNs promote animals to improve reward acquisition. Thus, our results demonstrate that the striatal direct and indirect pathways in TS have opposing functions in regulating threat management, allowing animals to adaptively overcome a threat via iMSN pathway while maintaining the threat estimate via dMSN pathway. The presence of this local opposing structure in TS provides fundamental insights into the design principle of the basal ganglia circuit; the threat-related controller (TS), while having its own opposing pathways within it, works in concert and competes, at a global scale, with reward-based behavioral controllers in other regions of the striatum, which are also equipped with direct and indirect pathways.

### Dynamical threat management and dopamine in TS

The present study found dynamical engagement of dopamine and striatal neurons in TS in threat management. Previous studies used various threatening stimuli such as a predator odor, a visual looming stimulus and a fictive predator to study avoidance behaviors (Choi and Kim, 2010; Cohen et al., 2006; Yilmaz and Meister, 2013). To study naturalistic threat management, recent studies established a semi-naturalistic foraging paradigm using a robot (robogator) (Amir et al., 2015; Choi and Kim, 2010). Similar to a robogator paradigm, we utilized a behavioral paradigm where animals need to balance a drive for water reward with a potential threat of an unknown object. We used an unknown object because this poses a common but difficult problem for animals; despite having limited knowledge of the novel object, animals must make judgements to estimate the potential for danger. Thus, while it is difficult to perfectly ensure safety, animals need to properly balance reward and threat to fulfill animals’ needs while avoiding harms.

In the monster paradigm, we found that mice displayed dynamical threat management with avoidance, prediction, and overcoming of threat manifesting at different stages. We focused on TS, because we previously found that dopamine in TS is important for avoidance of a potential threat including a novel object (Akiti et al., 2021; Menegas et al., 2018). Interestingly, phasic dopamine responses in TS are modulated by various parameters that are known to affect physical salience of external stimuli (Menegas et al., 2018), suggesting that an initial estimate of the threat level is shaped by physical salience conveyed by phasic TS dopamine responses (Akiti et al., 2021). In the present study, we systematically mapped dopamine activity pattern in the striatum, and found a correlational and causal relationships between physical salience, TS dopamine activity, and avoidance, further supporting this idea. Thus, while the actual threat outcome is unknown, animals may initially estimate the threat level using physical salience (i.e., size and movement) of the object, which is, in turn, used to learn threat prediction to determine future avoidance. Interestingly, animals showed avoidance at earlier and earlier locations in the arena, reminiscent of the incremental learning seen in the temporal difference (TD) learning (Sutton, 1988; Sutton and Barto, 1987, 1998), which is often used to model dopamine function (Montague et al., 1996; Schultz et al., 1997).

Although we observed dramatic effects of TS on threat management, TS had not been the focus of previous studies on threat-related behaviors. There may be two reasons. First, the location of TS might have caused previous researchers to miss TS. In humans, TS is an elongated structure aligned just beneath the hippocampus, thus it was proposed that TS was potentially categorized as hippocampus in functional imaging studies (Seger, 2013). On the other hand, TS in rodents is localized just above the amygdala, and thus it was potentially already ablated in some manipulation and recording/imaging studies. Second, the task design might be important. We found that manipulation of TS did not affect escape duration, but specifically affected hesitation in the pursuit of rewards. Thus, TS may play a role in more nuanced situations (rather than simple conditioning) where animals must estimate a “potential” threat, and/or under reward-threat conflicts. The monster paradigm, that we used in this study, might prove useful to study naturalistic threat management. Previous studies found that the basolateral and medial amygdala is critical for avoidance of a fictive predator (Amir et al., 2015; Choi and Kim, 2010; Miller et al., 2019). It will be important to study the relationship between TS and other established fear circuits (Amir et al., 2015; Branco and Redgrave, 2020; Choi and Kim, 2010; Hikosaka, 2010; LeDoux and Daw, 2018; Miller et al., 2019; Pare and Duvarci, 2012; Pereira and Moita, 2016) in threat management.

### Balancing avoidance and overcoming of a potential threat with dMSN and iMSN in TS

One of the major findings in this study is that the basal ganglia direct and indirect pathways may explain threat overcoming while maintaining threat estimation. Classically, the function of the direct and indirect pathways has been modeled as “scaling” and “focusing” (Alexander et al., 1990; Mink, 1996; Wichmann and DeLong, 1996) (Figure S6). With scaling, the strength of output signals is “scaled” by relative activity of direct and indirect pathways that compete each other (Figure S6, left). Thus, this model explains modulation of size of outputs. Different from scaling which models output size, “focusing” models choice of information. With focusing, while the direct pathway promotes a specific information flow, the indirect pathway inhibits other information, similar to the center surround suppression in the sensory systems. Thus, scaling works simply by the competition of two pathways, while focusing works by collaboration. Anatomically, in the former model, information from dMSNs and iMSNs in the same striatal area is focally integrated, for example in the substantia nigra (SN) or globus pallidus internal segment (GPi or EP) (Chen et al., 2021; Hikosaka et al., 1993), while in the latter, information from iMSNs spreads more by divergent projection through the indirect pathway and/or by differential output pathways from the globus pallidus external segment (GPe or GP) (Courtney, 2021; Hazrati and Parent, 1991; Kita, 2007; Shammah-Lagnado et al., 1996; Watabe-Uchida, 2019), although the precise integration mechanism remains to be determined. Functionally, these two models predict opposite phenotypes with manipulation of a neuron type. For example, since scaling is a competition between the two pathways, inhibition of iMSNs would increase the function of the direct pathway. On the other hand, since focusing indicates a collaborative function, inhibition of iMSNs would counter the function of the direct pathway, e.g. due to insufficient suppression of competing information/action.

In the present study, we found that dMSNs and iMSNs in TS functionally oppose each other, but at different stages, in threat management. dMSNs play a role in threat avoidance and prediction, while iMSNs play a role in overcoming threat (Figure S6, right). The relationship between these neurons resembles the idea of “scaling”; output strength (threat level) is scaled by relative activity of dMSNs and iMSNs. Since dMSNs directly inhibit SNL, while iMSNs indirectly disinhibit SNL, integration may partly happen in SNL. Thus, even if dMSNs and iMSNs show similar activity patterns overall, reflecting similar incoming inputs, TS may have the function of filtering out such concurrent activity by computing the difference. With this system, dMSNs and iMSNs can be specialized to learn from increases and decreases in dopamine levels, respectively. The parallel signaling of two types of information — threat estimates by dMSNs and overcoming threat by iMSNs — explains animals’ behaviors well since animals often need to overcome threats without erasing threat information. Further, parallel information flows could allow for independent top-down controls, depending on the situation. Notably, a previous study found that the prefrontal cortex (PFC) modulates TS signaling via GP (i.e. a part of the indirect pathway) to the thalamic reticular nucleus (TRN) so that animals can ignore distracting stimuli in an attention shifting task (Nakajima et al., 2019). Thus, iMSNs in TS may manipulate the sensory information to suppress attention toward it. Importantly, this suppression happened proactively, even before the distracting cue was presented (Nakajima et al., 2019). We also observed that iMSN activity started ramping up even before the monster started moving in successful trials. It is plausible that similar information (attention away from threat) from PFC to iMSN plays a role in overcoming threat under reward-threat conflict, in parallel with the indirect pathway competing with the direct pathway in SNL. The scaling mechanism using direct and indirect pathways allows for such flexible decision-making.

Our finding of scaling with dMSNs and iMSNs in TS indicates a critical modular organization in the basal ganglia and surprisingly similar architectures within each module. Opposing function through direct and indirect pathways has been widely observed in other striatal areas (Durieux et al., 2012; Kravitz et al., 2010; Kravitz et al., 2012; Nonomura et al., 2018). Previous studies found opposition of dMSNs and iMSNs along value axis (Hikida et al., 2010; Kravitz et al., 2012; Lobo et al., 2010; Stefanik et al., 2013), and along ipsilateral versus contralateral action values (Cruz et al., 2020; Lee and Sabatini, 2021; Tai et al., 2012). Thus, parallel signaling of opposing information within each striatal area may allow dynamical and flexible behavioral choice, as TS does for overcoming threat.

### Multi-agent reinforcement learning system specialized for outcome value and threat

Our finding that the direct and indirect pathway projection neurons in TS show opposing effects, albeit at different stages of threat management, was surprising, considering that value-based systems in the ventral and dorsal striatum already possess opposing circuits. For example, the ventral striatum is also important for threat avoidance, and direct and indirect pathway neurons have opposing function; dMSNs are important for reward approach and iMSNs for threat avoidance (Hikida et al., 2013; Yamaguchi et al., 2015). This is consistent with the observation that dopamine neurons in VTA represent integrated value of both appetitive and aversive events (Matsumoto et al., 2016). Similarly, in the amygdala, the same neurons represent both reward loss and aversive stimuli, suggesting that these neurons signal negative value of aversive stimuli (Paton et al., 2006; Shabel and Janak, 2009; Zhang et al., 2020). Consistent with these observations, classic machine learning models typically map various events or actions onto one dimensional value axis for action selection/learning (Dezfouli and Balleine, 2012; Samejima and Doya, 2007). However, our results revealed that outcome values and threats are represented along two separate axes. Thus, combining with the idea of the independent control of dMSN and iMSN, positive and negative value, and a threat estimate and overcoming the threat are all different information. Having specialists representing each type of information could make behaviors more flexible, as this would allow switching the controller of behaviors between these “specialists” depending on the situations, much similar to the multi-agent framework in recent reinforcement learning models (Ahilan and Dayan, 2019; Crites and Barto, 1998; Yang et al., 2019) .

How do threat and value information affect a behavioral choice? One possibility is that once TS computes a threat level, the threat information can be converted and sent to the value system so that the animal only needs to consider the integrated value for a behavioral choice. However, integration may not be the only strategy, since making decision solely on the basis of value predictions might not be flexible enough to balance reward and threat in the face of various situations. It is possible that decision about whether or not to engage with a potential threat are made based on two-dimensional information, weighing on each depending on situations, similar to the idea that emotion is regulated based on the valence and intensity axes (Anderson et al., 2003; Small et al., 2003).

Alternatively, TS may exert its function by “vetoing” other processing to prioritize the avoidance of a potential threat. TS is in a privileged position to receive and pass information faster than other striatal areas. TS is a part of the sensory basal ganglia, receiving visual and auditory information directly from the sensory cortices and thalamus (Hintiryan et al., 2016; Hunnicutt et al., 2016; Valjent and Gangarossa, 2021). Dopamine inputs to TS may be also faster (Fiorillo et al., 2013; Kim et al., 2015; Redgrave and Gurney, 2006). Notably, activity modulation in TS dopamine according to physical salience of sensory stimuli would be much easier than computation of value (Redgrave and Gurney, 2006). Further, SNL, the downstream of TS (Deniau et al., 1996; Gerfen, 1985), directly projects to the brain areas, such as periaqueductal gray and superior colliculus (Redgrave et al., 1992; Yasui et al., 1991), that can acutely induce avoidance (Assareh et al., 2016; Bandler et al., 1985; Branco and Redgrave, 2020; De Oca et al., 1998; Dean et al., 1989; DesJardin et al., 2013). Thus, for quick solutions to balance reward and threat, TS is probably indispensable while other striatal areas more accurately and deliberately compute overall value. In this sense, TS might function as a “first responder”, while other brain areas “calculate” for further actions.

At a more global circuit level, TS may be able to access information that value prediction areas may not. Importantly, TS indirectly projects back to cortices and thalamus that are important for sensory perception such as the temporal cortex and sensory parts of TRN (Middleton and Strick, 1996; Nakajima et al., 2019; Valjent and Gangarossa, 2021). This raises the possibility that TS may modify the sensory information even before other striatal areas can access the original information. Thus, TS may dominate in some situations when the TS outputs are strong enough to skew the sensory information.

Together, the present study provides insights into the design principle of the basal ganglia, pointing to a hierarchical, multi-agent organization whereby competing demands (e.g. reward and threat) are resolved in multiple steps involving competition within and across multiple agents. First, at a global scale, the striatum is parceled into multiple subsystems, e.g. specialized for value and threat prediction, defined by distinct dopamine signals and anatomical specializations. Second, each of these agents exerts its influence on behavior through functionally opposing circuits, i.e. direct and indirect pathways. Finally, behavioral outputs are controlled by sensibly weighing among these agents. While both value and threat prediction may be used for a behavioral choice, these subsystems compete against or coordinate with one another. In threatening situations, TS may take over due to its two unique anatomical features: because of direct sensory inputs and avoidance outputs, and because of its impact back onto sensory pathways. The modular architecture and the dominance of the threat system, indicated by the present and other studies, might be a sensible design of the brain which was shaped by the evolutionary process which would emphasize animals’ ultimate survival in uncertain environments.

## METHODS

### Animals

155 wild type mice, 12 tachykinin precursor 1 (Tac1)-cre (B6;129S-Tac1tm1.1(cre)Hze/J, Jackson Laboratory; RRID:IMSR JAX: 021877) heterozygous mice, 24 Tac1-cre;Ai14 (Rosa-CAG-LSL-tdTomato, Jackson Laboratory; RRID:IMSR JAX:007914) (Madisen et al., 2010) double heterozygous mice, 11 adenosine 2A receptor (Adora2A)-cre (B6.FVB(Cg)-Tg(Adora2a-cre)KG139Gsat/Mmucd, GENSAT; MGI:4361654) heterozygous mice, 24 Adora2A-cre;Ai14 double heterozygous mice, 6 vesicular glutamate transporter 2 (vGlut2)^lox/lox^ (Slc17a6tm1Lowl/J, Jackson Laboratory) heterozygous mice, 6 dopamine transporter (DAT)-cre (B6.SJL-Slc6a3tm1.1(cre)Bkmn/J, Jackson Laboratory; RRID:IMSR JAX:006660) (Bäckman et al., 2006);vGluT2^lox/lox^ double heterozygous mice, and 3 vGluT2-cre (B6J.129S6(FVB)-Slc17a6tm2(cre)Lowl/MwarJ, Jackson Laboratory) heterozygous mice, male and female, aged 8-20 weeks, were used. All mice were backcrossed with C57BL/6J (Jackson Laboratory). Animals were housed on a 12 hour dark/12 hour light cycle (dark from 07:00 to 19:00) and performed a task at the same time each day. Animals were group-housed (2-4 animals/cage) during training, and then single-housed after surgery. Some mice were water restricted for behavioral tests. In those cases, mice received water every day by experimenters and the body weights were kept >85% of their weights with freely available water. All procedures were performed in accordance with the National Institutes of Health Guide for the Care and Use of Laboratory Animals and approved by the Harvard Animal Care and Use Committee.

### Monster task

#### Apparatus

The monster apparatus (Figure 1A) was a long rectangular box (90 cm in length, 20 cm in width, 30 cm in height, white acrylic, product ID: 8505K755, McMaster-Carr, NJ) with ceiling. It was divided into two compartments with a door (height, 28 cm; width, 8 cm, clear red acrylic, product ID: 24163-02, INVENTABLES, IL), a smaller (12 cm-long) compartment (“shelter”) and a bigger (78 cm-long) compartment (“foraging arena”). To make the shelter dimmer (30 lux) than the foraging arena (100 lux), clear red acrylic (Product ID: 24163-02, INVENTABLES, IL) and transparent acrylic (product ID: 8536K162, McMaster-Carr, NJ) were used for ceiling of shelter and foraging arena, respectively. Both ceilings had a narrow slit (1 cm wide) in the center, to allow a patch cord attached to a mouse to follow animal’s movement in fluorometry experiments. A speaker (GHXamp, AliExpress, China) was attached on the wall at the far end of the foraging arena to present tones (see below). A door was opened and closed with a servo motor (product ID: 1143, Adafruit, NY). To detect animal’s position, infrared (IR) break beam sensors (product ID: 2168, Adafruit, NY) were installed in multiple locations on the wall (–8, –1, +1, +5, +10, +20, +30, +40 cm from the door; with a – denoting the homing side and a + denoting the foraging side). A waterspout was presented at trial start through a small hole on the floor at 40 cm from the door, and withdrawn at the end of the trial with a servo motor (product ID: 169, Adafruit, NY). Animal’s licking was detected with a touch sensor (product ID: 1982, Adafruit, NY) attached to the waterspout. All electronics were controlled by Teensy 3.2 (SparkFun Electronics, CO) and Python software (https://www.python.org/).

#### Monster

In some sessions, an object (18 cm in height, 18 cm in width, 15 cm in depth, Jurassic World Velociraptor Blue 1/2 Mask, Rubies, NY) (“monster”) was placed at the far end of the foraging arena, facing the homing arena. A monster was attached to a gear rack (30 cm long, product ID: 7854K15, McMaster-Carr, NJ), which penetrated the wall at the far end of the foraging arena through a narrow slit (1cm wide). Monster movement was controlled with a servo motor (product ID: CPM-MCVC-2310S-RLN, TEKNIC, NY), connected to a gear (product ID: 57655K54, McMaster-Carr, NJ) that is located outside of the arena. A “monster territory” was defined as the far side of a foraging arena (30 cm or further from the door). When a mouse entered the monster territory (sensed with IR beam break at 30 cm from the door), a monster started to move forward (10 cm at 20 cm/s). After the forward movement, the monster stayed for 500 ms during which a loud complex tone (150 dB, Godzilla Sounds, SoundBible.com, https://soundbible.com/tags-godzilla.htm) was presented, and then returned to its original position (20 cm/s). The back and forth movement with tone was repeated until the mouse returned to the shelter. To test the effects of monster movement on avoidance behavior, a monster was placed in the same manner but the motor was turned off (“static monster”) (Figure 2E). To test effects of size of the monster on avoidance behavior, smaller objects (“medium monster”, 10 cm in height, 5 cm in width, 7 cm in depth, Dinosaur Toy Untamed T-Rex, Shenzhen ZCT Technology, China; “small monster”, 3 cm in height, 2 cm in width, 6 cm in depth, Mini Dinosaur Play Set, Zooawa, China) were used (Figure 5A-D).

#### Training

On the first day of habituation, mice were handled by an experimenter for 10 min. Water restriction was started and continued until the last day of the behavioral testing. On the following day, mice were placed in the shelter for 30 min with droplets of water and food on the floor to acclimate to the area. Then, 3 days of training sessions started. A mouse was gently introduced in the shelter with the door closed. A trial was initiated with the door opening. The entry to the foraging arena was detected when the mouse broke the IR beam at 5 cm from the door. During a trial, a mouse was allowed to freely explore the foraging arena. When a mouse licked a waterspout, a drop of water (10 μl) was delivered. Water reward was available only once per trial so that a mouse was not rewarded even if it continued to lick the waterspout. When a mouse did not enter the foraging arena for 180 sec, the door was closed and the trial was ended. Between sessions, the arena was thoroughly cleaned, with the base of the arena was wiped down with 70% ethanol. 10 trials were run per session per day.

#### Behavioral tests

Following the 3 days of training, animal behaviors were tested for 7 days (4 days with no monster and 3 days with monster, interleaved in an alternating manner). Control (no monster) sessions were exactly the same as training sessions. Monster sessions were the same as control sessions except for the presence of a monster. The monster moved when a mouse entered the monster territory (see the above, “Monster”).

Failure of reward acquisition was defined as any trial without reward acquisition (a contact to the water spout). A failure rate for reward acquisition for each trial was calculated as the fraction of animals that failed, and SEM was calculated using binomial distribution. A failure rate for each session or multiple sessions was calculated as the fraction of trials that each animal failed, and then mean and SEM in all animals were obtained. Failure before monster movement was defined as any trial without entry to the monster territory. A rate of failure before monster movement was calculated as the fraction of animals that did not enter the monster territory (each trial data) or as the fraction of trials when each animal did not enter the monster territory (session data). Failure after monster movement was defined as any trial when animals enter the monster territory but did not acquire reward. A rate of failure after monster movement was calculated as the fraction of animals that did not acquire water in animals that entered a monster territory (each trial data) or as the fraction of trials that did not acquire water in trials when each animal entered a monster territory (session data).

#### Drug infusion

To inject a dopamine transporter (DAT) inhibitor (GBR12909, D052, Sigma Aldrich, MO, 5 mg/ml in distilled water with 5% dimethyl sulfoxide, 67-68-5, Sigma Aldrich, MO) or vehicle (distilled water with 5% dimethyl sulfoxide), we followed an existing protocol (Mazei et al., 2002; Menegas et al., 2018). The cannula plug (see Surgical procedures) was removed and replaced with an infusion needle (4.2 mm long, C317I/SPC, P1 Technologies, VA). A solution (300 nl/side with 200 nl/min flow rate) was infused with a syringe pump (70-4501, Harvard Apparatus, MA), which was connected to the infusion needle via a polyethylene tube (50 cm long, C313CT/PKG, P1 Technologies, VA). Following the injection, the infusion needle was left in the brain for 5 min. Then, the infusion needle was removed and replaced with the cannula plug. For validation of effects of DAT inhibitor (Figure 5C), solution was unilaterally infused in a head-fixed animal. For behavioral tests, solution was bilaterally infused while an animal was freely moving in the home cage (Figure 5D).

### Head-fixed task

#### Tone-water test for dopamine functional mapping

We followed an existing protocol for the head-fixed tone-water test (Menegas et al., 2018). After recovery from surgery, mice were handled for 10 min and water restriction was started and continued until the final day of behavioral testing. Then, mice were habituated to being head-fixed for 3 days. During these days, mice were head-fixed for 5–10 min and given water at random intervals (exponential distribution between 10–20 sec, average 13 sec). After habituation, dopamine sensor signals were recorded for 1 session while mice performed in tone-water tests (see “Fluorometry (photometry) recording”); 3 intensities of 5 kHz pure tone (50 dB, 75 dB, and 100 dB) and 3 sizes of water (1 μl, 3 μl, and 10 μl) were presented in pseudo-random order. Each session consisted of 120 trials.

#### Tone test for validation of DAT inhibitor

After recovery from surgery, mice were handled and habituated with the head-fixed preparation as described above. Then, dopamine sensor signals were recorded while mice performed in a tone test (pre-test) (see “Fluorometry (photometry) recording”); 3 intensities of a complex tone (50, 75 and 100 dB) were presented for 1 sec in a pseudo-random order. Complex tones (Incredible Free Sound Effects, Mixkit. Co., https://mixkit.co/free-sound-effects/) were played with LabView software (National Instruments, TX). After the test, mice were kept head-fixed and either DAT inhibitor or vehicle was injected into the TS (see Drug infusion). During the injection periods, the laser path was covered to prevent photobleaching. After the injection, dopamine sensor signals were recorded while mice were presented with the same tones used in the pre-test (post-test). Each session (before or after drug infusion) consisted of 24 trials. Dopamine sensor signals were normalized with mean responses to 75 dB tone (0-1 s after tone onset) in pre-test.

### AAV construct

To make DNA construct for AAV to express diphtheria toxin subunit A (dtA), PGKdtabpA (gift from Philippe Soriano; Addgene, #13440) (Soriano, 1997) was cleaved with NcoI and SacI, blunted and was subcloned into pAAV-CA-FLEX (Addgene, #38042) (Watabe-Uchida et al., 2012) cleaved with EcoRV to obtain pAAV-CAG-FLEX-dTA. AAV was produced at UNC vector core. The construct will be deposited at Addgene.

### Surgical procedures

All surgeries were performed under aseptic conditions with animals anesthetized with isoflurane (1–2% at 0.5–1.0 l/min). Analgesia was administered pre- (buprenorphine, 0.1 mg/kg, I.P) and post-operatively (ketoprofen, 5 mg/kg, I.P). We used the following coordinates to target injections and/or implants for TS: Bregma: –1.5 mm, Lateral: +3.2 mm, Depth: –2.4 mm, tDLS/aTS: Bregma: –1.0 mm, Lateral: +3.2 mm, Depth: –2.4 mm, cDLS: Bregma: –0.5 mm, Lateral: +3.0 mm, Depth: –2.4 mm, and SNL: Bregma: –3.5 mm, Lateral: +1.9 mm, Depth: –3.8 mm (relative to dura) (Paxinos and Franklin, 2019).

#### Ablation of dopamine neurons

To bilaterally ablate dopamine neurons projecting to the striatum, we followed an existing protocol (Akiti et al., 2021; Menegas et al., 2018; Thiele et al., 2012). The following solution was injected (I.P.) to animals at 10 ml/kg: 28.5 mg desipramine (D3900, Sigma-Aldrich, MO) and 6.2 mg pargyline (P8013, Sigma-Aldrich, MO) in 10 ml water. This was given to prevent uptake of 6-hydroxydopamine (6-OHDA) by noradrenaline neurons and to increase the selectivity of uptake by dopamine neurons. After injection, mice were anesthetized as described above. We then prepared a solution of 10 mg/ml 6-OHDA (H116, Sigma-Aldrich, MO) dissolved in 0.2% ascorbic acid (1043003, Sigma-Aldrich, MO) in saline (0.9% NaCL; PHR1008, Sigma-Aldrich, MO). The ascorbic acid in this solution helps prevent 6-OHDA from breaking down. Control animals were injected with 0.2% ascorbic acid solution (vehicle). To further prevent 6-OHDA from breaking down, we kept the solution on ice, wrapped in aluminum foil, and it was used within three hours of preparation. If the solution turned brown (indicating that 6-OHDA has broken down), it was discarded, and fresh solution was made. 6-OHDA (or vehicle) was injected bilaterally into cDLS, tDLS/aTS, or TS (200 nl per side). Mice were given 1 week resting to recover and to allow for sufficient cell death to occur. Control animals were pooled for Figure 1 and 2.

#### Guide-cannula implantation surgical procedure

To inject a dopamine transporter (DAT) inhibitor into the TS, we bilaterally (for behavioral tests) or unilaterally (for system verification) implanted a stainless guide cannula (4 mm long, 23 GA, C317G/SPC, P1 Technologies, VA). We slowly lowered the cannula into the TS, one side at a time. Once cannula was lowered, we attached it to the skull with black Ortho-Jet dental adhesive (Ortho-Jet, Lang Dental, IL). After waiting 15 min for the dental adhesive to dry, we applied a very small amount of rapid-curing epoxy (A00254, Devcon, MA) to attach the cannula even more firmly to the underlying adhesive. After waiting 15 min for the epoxy to dry, a cannula plug (4.2 mm long, 30 GA, C317DC/SPC, P1 Technologies, VA) was inserted to prevent tissue growth in the cannula. Mice were given 1 week of recovery time to rest following the procedure.

#### Fluorometry (photometry) surgical procedure

To express GRAB_DA2m_ (Sun et al., 2020), we unilaterally injected 300 nl of mixed (1:1) virus solution; AAV9-Syn-GRAB_DA2m_ (5.0×10^13^ particles/ml, Vigene Bioscience, MD) and AAV5- CAG-tdTomato (4.3×10^12^ particles/ml, UNC Vector Core, NC) into the striatum. For specific expression of GCaMP7f in dMSNs and iMSNs, we unilaterally injected 300 nl of mixed (1:1) virus solution; AAV9-Syn-FLEX-GCaMP7f (Dana et al., 2019) (2.8×10^13^ particles/ml, catalog#: 104492, Addgene, MA) and AAV5-CAG-FLEX-tdTomato (7.8×10^12^ particles/ml, UNC Vector Core, NC) into TS in Tac1- or Adora2A-cre mice. Virus injection lasted around 20 minutes, after which the injection pipette was slowly removed over the course of several minutes to prevent damage to the tissue. We also implanted an optic fiber (400 µm diameter, Doric Lenses, Canada) into the virus injection site. To do this, we first slowly lowered an optical fiber into the striatum. Once the fiber was lowered, we first attached it to the skull with UV-curing epoxy (NOA81, Thorlabs, NJ), and then a layer of rapid-curing epoxy to attach the optical fiber even more firmly to the underlying glue. After waiting 15 minutes for this to dry, we applied a black dental adhesive (Ortho-Jet, Lang Dental, IL). We used a zirconia ferrule (ZF_FLT, Doric Lenses, Canada) for a corresponding patch cord (SMA-MF, Doric Lenses, Canada) in head-fixed experiments and a magnetic fiber cannula (SMR_FLT, Doric Lenses, Canada) for a patch cord (SMA-SMC, Doric Lenses, Canada) in the freely-moving experiments. After waiting 15 minutes for the dental adhesive to dry, the surgery was complete.

#### Other AAV surgical procedure

To specifically ablate dMSNs or iMSNs in the striatum, AAV8-CAG-FLEX-diphtheria toxin subunit A (dtA, 2.6×10^12^ particles/ml) was bilaterally injected (200 nl each) into cDLS, tDLS/aTS, or TS in Tac1-cre/tdTomato mice, and in the TS in Adora2A-cre/tdTomato mice. Saline was bilaterally injected in TS in control mice. Injection procedures are the same as described in Fluorometry surgical procedures. Mice were given 2 weeks resting to recover and to allow cell death.

To visualize vGluT2-positive neurons in the SNL, 300 nl of AAV5-CAG-FLEX-GFP (4.3×10^12^ particles/ml, UNC Vector Core, NC) was unilaterally injected into SNL in vGluT2-cre mice. Injection procedures are the same as described in Fluorometry surgical procedures. Injection procedures are the same as described in Fluorometry surgical procedures. Histology was performed after 2 weeks from the surgery.

### Histology and immunohistochemistry

Histology was conducted in the same manner as previously reported (Akiti et al., 2021; Menegas et al., 2018). Mice were perfused using 4% paraformaldehyde, then brains were sliced into 100 µm thick coronal sections using a vibratome (Leica, Germany) and stored in PBS. To visualize dopamine axons in the striatum and dopamine cell bodies in the midbrain, brain sections were incubated with rabbit anti-tyrosine hydroxylase antibodies (TH; AB152, MilliporeSigma, MO) at 4°C overnight and then with fluorescent secondary antibodies (A-11012, Thermo Fisher Scientific, MA) at 4°C overnight. Slices were then mounted in 4’,6-diamidino-2-phenylindole (DAPI)-containing anti-fade solution (VECTASHIELD anti-fade mounting medium, H-1200, Vector Laboratories, CA) and imaged with Zeiss Axio Scan Z1 slide scanner fluorescence microscope (Zeiss, Germany).

### Fluorometry (photometry) recording

Fluorometry recording was performed as previously reported (Akiti et al., 2021; Tsutsui-Kimura et al., 2020). We used an optic fiber to stably access deep brain regions and interface with a flexible patch cord on the skull. The patch cord simultaneously delivers excitation light (473 nm, Laserglow Technologies, Canada; 561 nm, Opto Engine LLC, UT) and collects dopamine sensor/GCaMP and tdTomato fluorescence emissions. Activity-dependent fluorescence emitted by cells in the vicinity of the implanted fiber’s tip (NA=0.48) was spectrally separated from the excitation light using a dichroic, passed through a single band filter, and focused on a photodetector connected to a current preamplifier (SR570, Stanford Research Systems, CA). During photometry recording, optic fibers on the animal’s skull were connected to a magnetic (400 µm diameter, NA 0.48, 3 m long, SMA-SMC, Doric Lenses, Canada) or zirconia (SMA-MF, Doric Lenses, Canada) patch cord for freely moving or head-fixed experiments, respectively. The emitted light was then filtered using a 493/574 nm beam splitter (Semrock, NY), followed by a 500 ± 20 nm (Chroma, VT) and 661 ± 20 nm (Semrock, NY) bandpass filter, and collected by a photodetector (FDS10 X 10 silicone photodiode, Thorlabs, NJ) which is connected to a current preamplifier (SR570, Stanford Research Systems, CA). This preamplifier outputs a voltage signal which was collected by a data acquisition board (NIDAQ, National Instruments, TX) and custom software written in Labview (National Instruments, TX). Lasers were turned on at least 30 minutes prior to recording to allow them to stabilize. Before each recording session, laser power and amplifier settings were adjusted. After each recording session, collected light intensity was measured from the patch cord using a photometer. Light intensity fell within a range of 50-200 μW across animals and days.

#### Signal analysis

DA sensor or GCaMP (green) and tdTomato (red) signals were collected as voltage measurements from current pre-amplifiers (SR570, Stanford Research Systems, CA). Green and red signals were cleaned by removing 60 Hz noise with band-stop, finite impulse response (FIR) filter at 58-62 Hz and smoothing with a moving average of signals in 50 ms. The global change within a session was normalized using a moving median of 100 s. Then, the correlation between green and red signals was examined by linear regression. If the correlation was significant (p<0.05), the fitted red signals were subtracted from green signals. Responses aligned at a behavioral event were calculated by subtracting the average baseline activity (-2 s to -0.2 s before trial start).

We built a regularized linear regression to fit cosine kernels (Parker et al., 2016; Tsutsui-Kimura et al., 2020) (width of 500 ms, interval of 100 ms) to the activity of dopamine axons in each animal. We used down-sampled (every 20 ms) responses for the model fitting. We used two different time points to lock kernels: water onset (‘water’) and monster movement onset (‘monster’). Both kernels span -5s to 12s from the event start. All the kernels were fitted to responses using linear regression with Elastic net regularization (α=0.75) with 10-fold cross validation. The regularization coefficient lambda was chosen so that cross-validation error was minimum plus one standard deviation. A percent explained by a model was expressed as the reduction of the variance in the residual responses compared to the original responses. Contributions of each component in the model were measured by reduction of the deviance compared to a reduced model excluding the component.

### Statistical analyses

The experimenters were blinded to the treatments of mice in ablation studies until completion of behavioral analyses. The number of animals used for ablation studies (dopamine neurons and dMSN, n=6 animals) was determined by a power analysis using a pilot experiment (t-test, 6 animals for control and 6 animals for dopamine neuron ablation, data not included because the apparatus was slightly different) to be able to detect a significant difference in the failure rate for water acquisition compared to controls at 90% confidence level. The number of animals used for ablation studies (iMSN, n=12 animals) was determined by a power analysis using a pilot experiment (t-test, 6 animals for iMSN ablation and 18 animals for pooled controls) to be able to detect a significant difference in improvement of the failure rate at 90% confidence level. Data analysis was performed using custom software written in MATLAB (MathWorks, Natick, MA). All error bars in the figures are SEM unless notification was given. In boxplots, the edges of the boxes are the 25^th^ and 75^th^ percentiles, and the whiskers extend to the most extreme data points not considered outliers.

### Data Availability

All data (both behavioral and fluorometry) will be deposited.

## ACKNOWLEDGEMENTS

We thank Michael Bukwich, Malcolm Campbell, Adam Lowet, Sara Matias and all Uchida lab members for discussion, Brett Graham and Edward Soucy for monster apparatus, Ryunosuke Amo for AAV-CAG-FLEX-dtA, Takahiro Yamaguchi for a LabView code, and William Menegas for fluorometry setup. This work was supported by NIH BRAIN Initiative (U19NS113201, NU; and R01NS108740, NU), National Institute of Mental Health (R01MH125162, MW-U), Simons Collaboration on the Global Brain (NU), Bipolar Disorder Seed Grant Program (NU), and Japan Society for the Promotion of Science (IT-K).

## AUTHOR CONTRIBUTIONS

IT-K, NU, and MW-U initiated the project. IT-K performed experiments. IT-K and MW-U analyzed data. IT-K and MW-U wrote the paper and IT-K, NU, and MW-U edited the paper. MW-U supervised the project.

---

**Figure S1.**
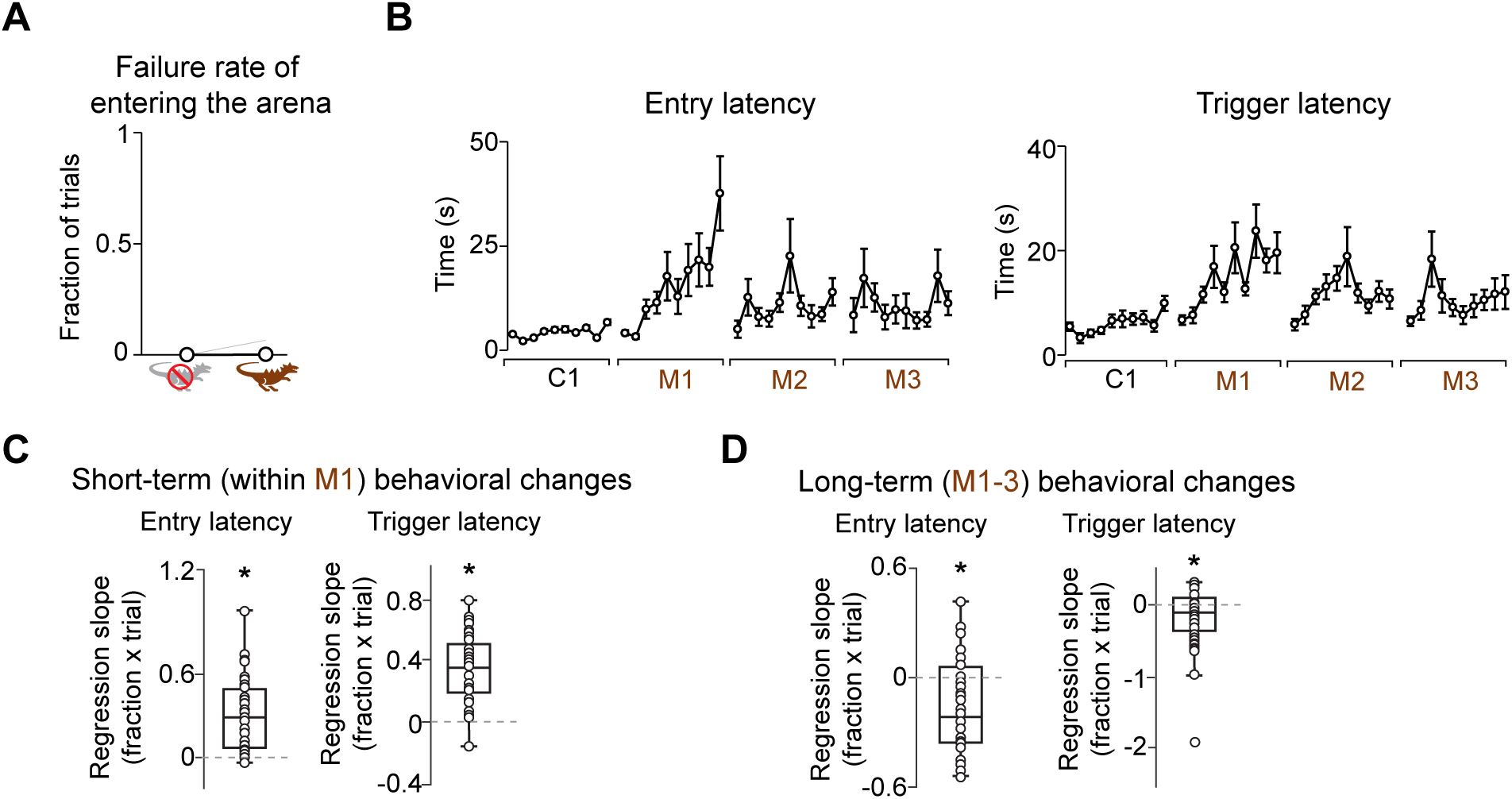
Mice hesitate in the presence of a monster. (**A**) The average failure rate of entering the foraging arena. Almost all animal (23/24) entered the arena even in the presence of the monster (monster sessions vs control sessions, p=0.33, paired t-test). (**B**) Time-course of the latency from door opening to entry to the foraging arena (entry latency, left) and the latency from door opening to entry to the monster territory (trigger latency, right). (**C**) Regression coefficients of entry latency (left, p=5.7×10^-6^, t-test) and of trigger latency (right, p=7.0×10^-8^, t-test) with trial number in M1 for each mouse. Entry latency and trigger latency gradually increased over trials within M1. (**D**) Regression coefficients of entry latency (left, p=0.012, paired t-test) and trigger latency (right, p=0.043, paired t-test) with trial number in M1 to M3 in each animal. Entry latency and trigger latency became gradually shorter over the multiple monster sessions.

**Figure S2.**
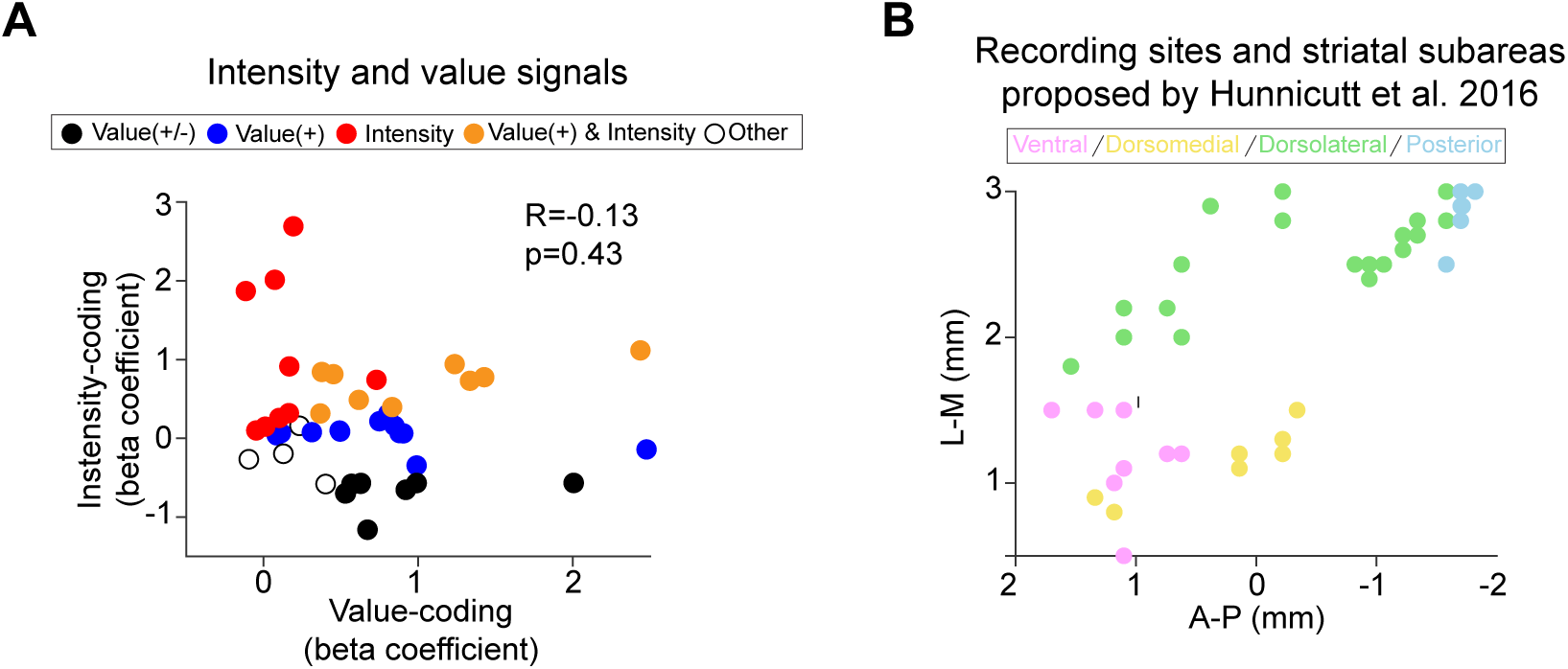
Value-modulation vs intensity-modulation in dopamine responses in the striatum. (**A**) Comparison of dopamine activity modulation with reward amount and tone intensity. Linear regression beta coefficients of dopamine activity with reward amount and beta coefficients of dopamine activity with tone intensity showed negative correlation (R=-0.13, p=0.43, Pearson’s correlation coefficient). (**B**) Location of dopamine recordings (Figure 3) and relation to striatal subareas categorized by cortico-striatal inputs (Hunnicutt et al., 2016).

**Figure S3.**
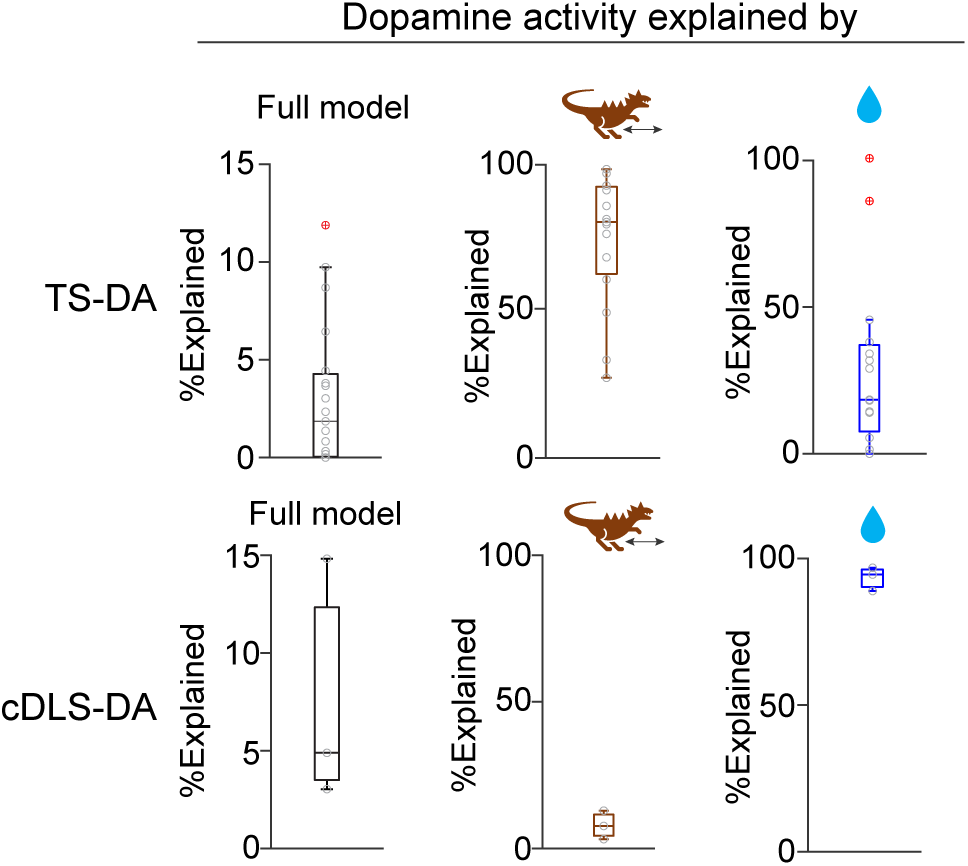
Kernel regression models of dopamine signals in a monster paradigm. Contribution of each component in the kernel regression models of TS-DA (top) and cDLS-DA (bottom) (see Methods).

**Figure S4.**
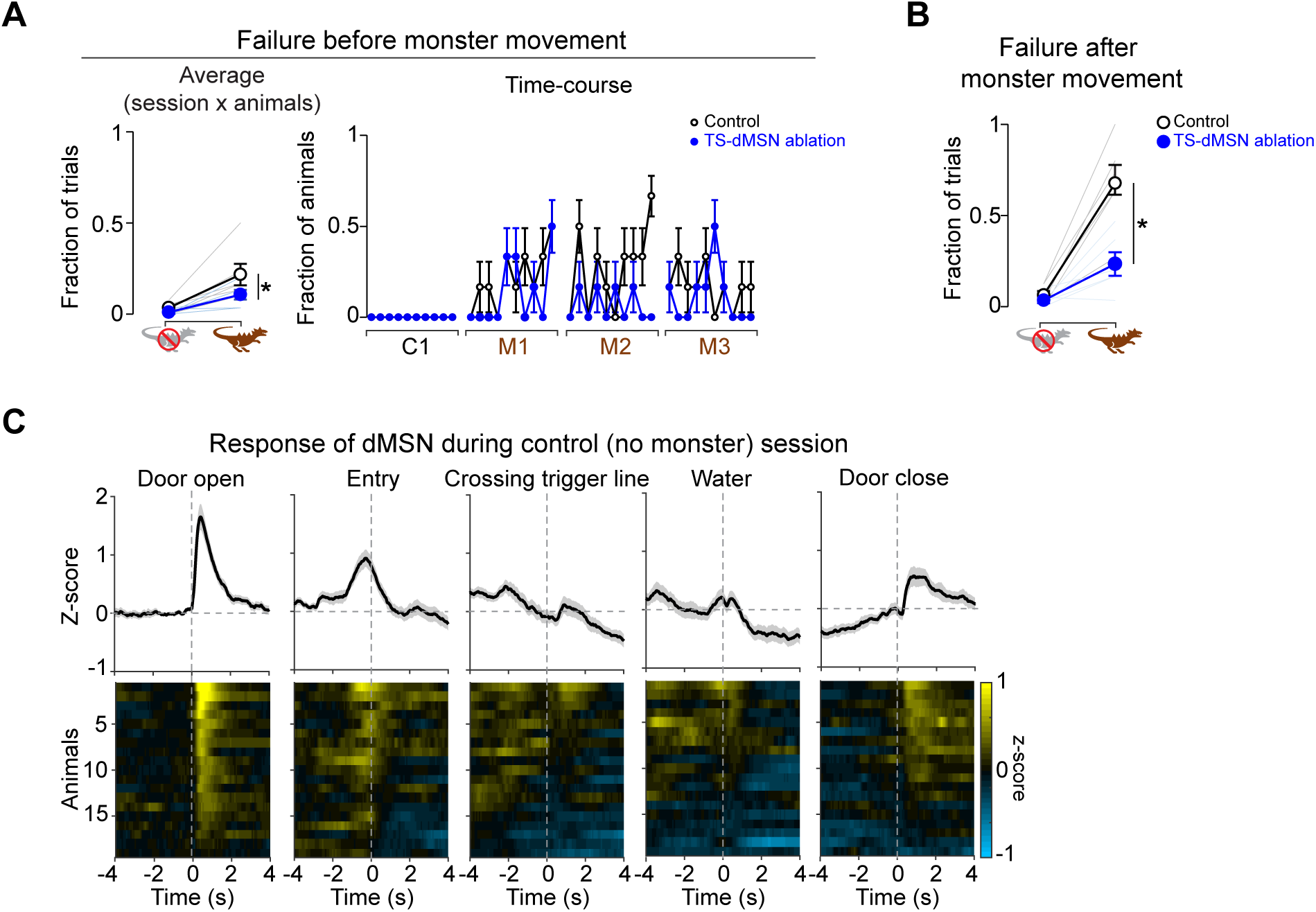
TS-dMSN ablation and recording. (**A**) Left, the average rate of failure before monster movement was lower in TS-dMSN ablation mice than control mice (p=0.045, t-test). Right, the time-course of the rate of failure before monster movement. Error bars, SEM (binomial). (**B**) The average rate of failure after monster movement was lower in TS-dMSN ablation mice than the control mice (p=1.2×10^-3^, t-test). (**C**) dMSN activity in the control session with no monster (mean ± SEM).

**Figure S5.**
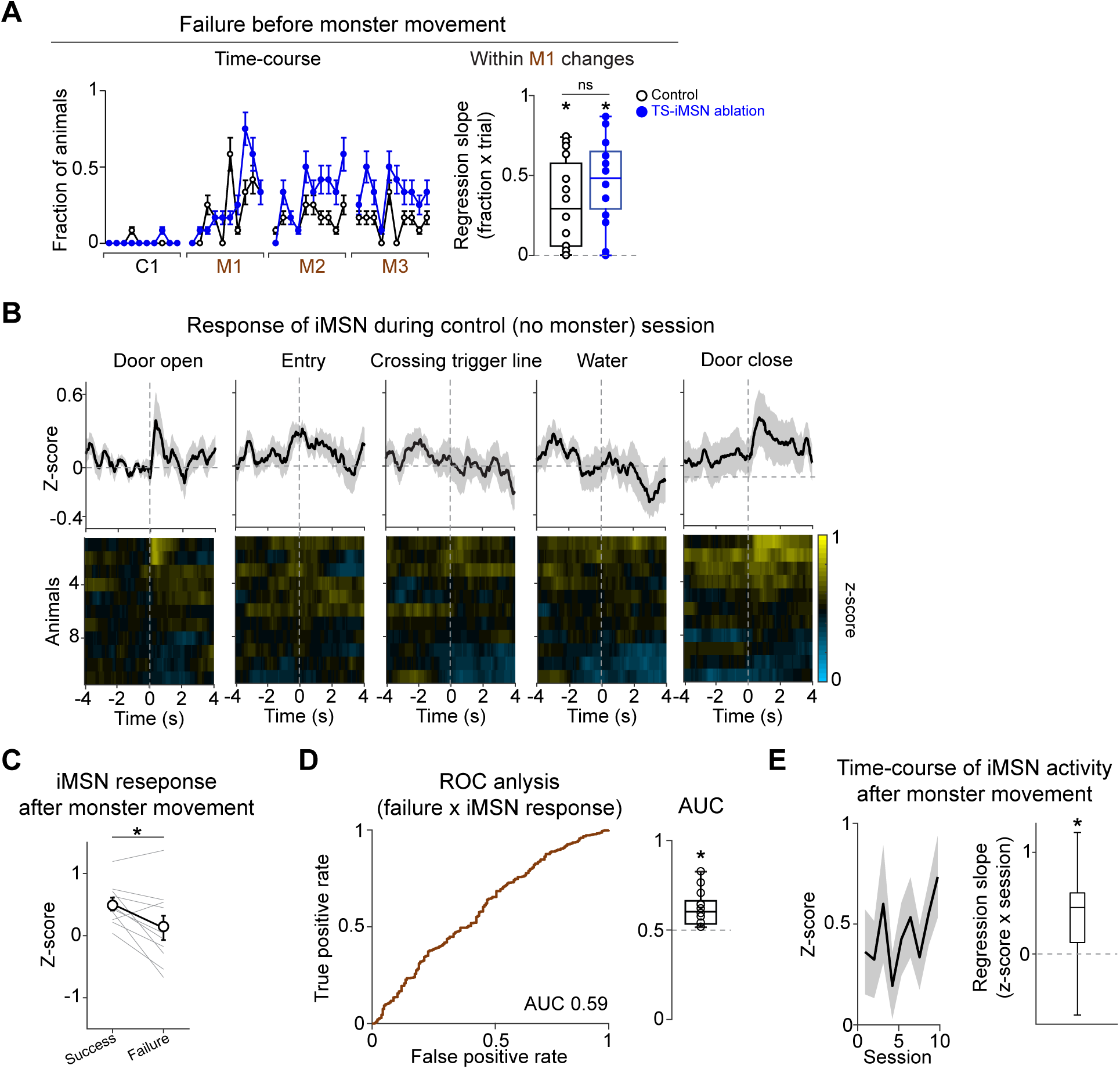
TS-iMSN ablation and recording. (A) Left, the time-course of the rate of failure before monster movement. Error bars, SEM (binomial). Right, the rate of failure before monster movement was gradually increased over trials in M1 in both control (left, p=1.3×10^-4^, t-test) and TS-iMSN ablation (left, p=1.8×10^-3^, t-test) mice (control vs ablation, p=0.30, t-test). (**B**) iMSN activity in the control session with no monster (mean ± SEM). (**C**) iMSN responses to monster movement were higher in success trials than in failure trials (p=0.047, paired t-test). (**D**) Left, ROC curves evaluating performance of the iMSN activity after monster movement (bottom, AUC=0.59) against success/failure outcome. Right, AUC for each animal (0.61 ± 0.09, mean ± STD; AUC vs 0.5, p=4.3×10^-3^, t-test). (**E**) iMSN responses to monster movement increased across sessions (left, linear regression of average iMSN responses with session number, coefficient beta=0.033, p=0.13; right, beta coefficients of each animal, p=0.035, t-test).

**Figure S6.**
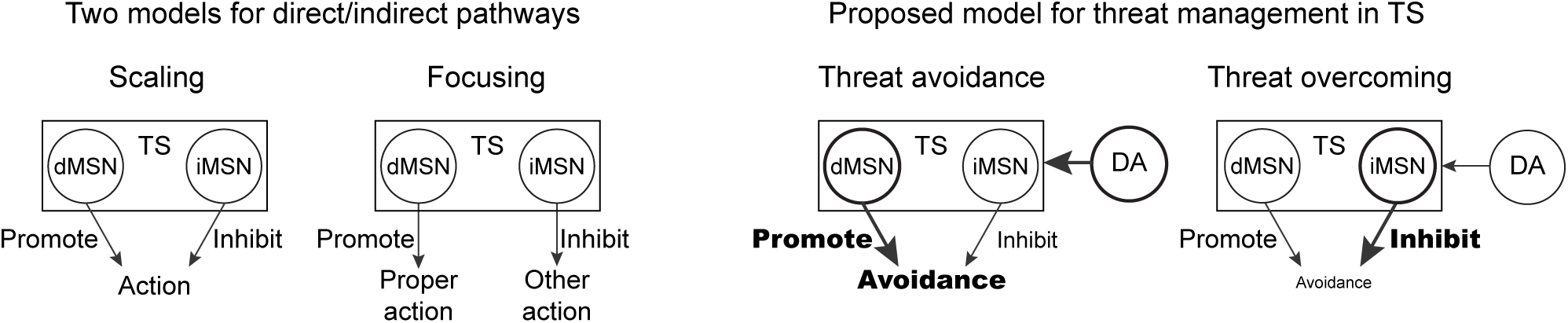
Differential roles of dopamine and direct and indirect pathway neurons in TS in threat management. Left, classical models for striatal direct/indirect pathways; “scaling” and “focusing” (Alexander et al., 1990; Mink, 1996; Wichmann and DeLong, 1996). Right, we demonstrated that the TS direct pathway neurons promote avoidance and the indirect pathway neurons inhibit avoidance, which resembles “scaling” model on the left.

